# Protein Language Model Decoys for Target Decoy Competition in Proteomics: Quality Assessment and Benchmarks

**DOI:** 10.64898/2026.03.27.714819

**Authors:** Grigory Reznikov, Fabrice Kusters, Majid Mohammadi, Henk W.P. van den Toorn, Pavel Sinitcyn

## Abstract

Large-scale proteomics relies heavily on target–decoy competition for false discovery rate estimation in peptide identification, and the performance of this strategy depends strongly on the design of the decoy database. Classical generators such as reversal and shuffling remain widely used. Here, we introduce the first protein language model-based (PLM) decoy generation for peptide identification and benchmark it against classical strategies. We evaluate these approaches using three complementary quality-control layers: sequence-based separability, search-engine-agnostic spectral-space diagnostics, and end-to-end mass spectrometry benchmarks, including pipelines with rescoring. Across these analyses, PLM-based decoys are harder for sequence-only neural networks to distinguish than most classical generators, suggesting fewer obvious sequence-level artifacts. However, this signal is only weakly informative for search performance. Spectral diagnostics further show that short peptides occupy a particularly crowded target–decoy space and are therefore especially prone to local collisions across all generators. In full search pipelines, reverse decoys remain a strong baseline, and current PLM-based generators do not yet provide a clear overall advantage. We therefore view PLM-based decoys not as universal replacements for reverse decoys, but as tunable tools for benchmarking, diagnostics, stress testing, and future adaptive decoy optimization, with increasing value as search models become more expressive.

## Introduction

Target–decoy competition (TDC) remains the standard approach for false discovery rate (FDR) estimation in most proteomics pipelines. ^1–3^In the standard view, decoy hits approximate false target hits and thereby provide a practical estimate of the null score distribution. In most workflows, decoys are generated with simple sequence transformations such as reversal or shuffling, which are attractive because they are fast, easy to implement, and usually work well in practice. However, modern search pipelines increasingly rely on machine learning-based (ML) scoring and rescoring techniques. ^4–8^This raises a concern that overly easy decoys may expose shortcut signals tied to how the decoys were constructed, allowing a model to separate targets from decoys without relying on peptide–spectrum match (PSM) evidence. In that regime, estimated FDR can become optimistic and accepted target sets can contain more false positives than expected. ^9,10^

One possible response is to replace hand-crafted decoys with learned ones. Protein language models (PLMs) are self-supervised models trained on large collections of protein sequences, and models such as Evolutionary Scale Modeling 2 (ESM2) have shown that PLMs can capture substantial statistical structure in protein sequence space. ^11,12^This makes PLMs a promising candidate for decoy generation, as they may produce sequences that appear less artificial than simple reversed or shuffled peptides, an approach that, to the best of our knowledge, has not previously been explored. At the same time, it is not obvious that sequence realism alone is the property that matters most for TDC and, consequently, for proteomics search results.

To make this question more precise, we revisit the conditions that adequate decoys should satisfy in TDC. First, on null spectra (i.e. spectra with no true target match), targets and decoys should win with equal probability. ^13^Second, on spectra generated by true peptides, decoys should not beat the corresponding target. ^14^These requirements are natural, but they are not determined by the decoy generator alone. Instead, they depend jointly on the search engine, the data, and the target and decoy databases. As a result, decoy adequacy is difficult to verify directly and even harder to optimize in a model-agnostic way.

We therefore study decoy quality through three complementary experiments (Figure 1A). The first asks whether targets and decoys can already be distinguished from sequence alone. Although sequence-only separability is not itself a criterion of TDC validity, it serves as a useful screen for construction-related information leakage. The second provides a search-engine-agnostic but mass-spectrometry-relevant view by representing peptides with *in silico* predicted spectra and measuring distances in cosine space. The third evaluates decoys end-to-end in real search pipelines on experimental datasets. To anchor these diagnostics, we also introduce two stress-test generators: a deliberately “easy” `Random` and a deliberately “hard” `Isobaric` generators.

**Figure 1:**
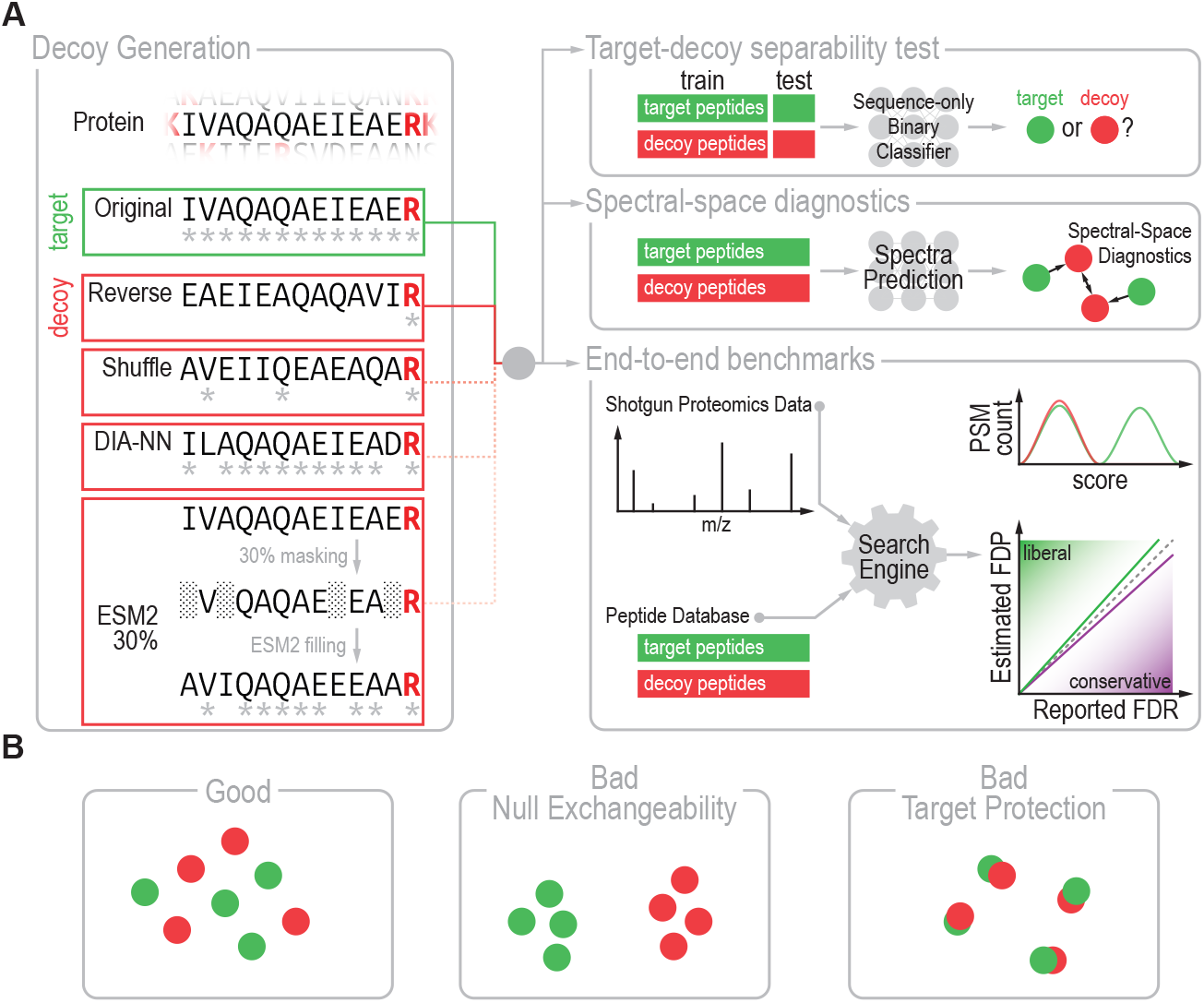
Overview of the study workflow. **(A)** Study workflow, including the decoy generation step and three evaluation approaches: target–decoy separability testing, spectral-space diagnostics, and end-to-end benchmarking. **(B)** Schematic illustration of spectral-space arrangements for three decoy scenarios: good, where targets and decoys are well matched both globally and locally; bad, with violation of null exchangeability, where targets and decoys are globally separable; and bad, with violation of target protection, where target and decoy peptides are too close to each other.

Taken together, our results give a consistent picture. Reverse decoys remain a strong practical baseline under current search settings. ESM-based generators are less sequence-separable, but this does not translate into a statistically significant gain in end-to-end identification performance. At the same time, the diagnostics reveal systematic structure in target–decoy space: short peptides are intrinsically more vulnerable, and close target–decoy collisions in reverse databases are often driven by *I*↔ *L* equivalence or local residue swaps. We therefore frame PLM-based decoys not as an outright replacement for classical decoys today, but as a tunable family for rare use cases, diagnostics, stress testing, and for probing how future, more expressive search models interact with decoy construction.

We release the generator and evaluation pipeline as open-source software for benchmarking decoy-generation methods and probing search-engine failure modes. Code is available at https://github.com/SinitcynLab/DecoyGeneration.

## Methods

### Decoy generators

Classical sequence-decoy strategies were implemented as lightweight transformations of target peptides. In protease-aware settings, digestion-defining boundary residues were preserved, and all other position were treated as mutable.

#### Reverse

Sequence order was reversed while preserving the digestion-defining boundary residues. For a tryptic peptide, the C-terminal K/R residue was kept fixed.

#### Sage

Internal residues were reversed while the first and last amino acids were kept fixed, following the algorithm used in the Sage search engine. ^15^

#### Shuffle

Residues were randomly permuted under the same digestion-aware constraints.

#### DIA-NN

Local mutations were introduced at the first residue and near the peptide C terminus while preserving tryptic boundary residues, following the DIA-NN-style heuristic. ^6^The replacement rule is defined in the Table S2. This decoy method was used in DIA-NN up to version 1.7.12 according to the last publicly available source code, and up to version 1.8.1 according to the DIA-NN author.

#### Random

Each target peptide was replaced with a random 15-residue peptide.

#### Isobaric

Near-isobaric edits were applied using operations such as *I* ↔ *L, GG* ↔ *N*, and local *XY* ↔ *Y X* swaps.

#### ESM2-based generators

PLM-generated decoys were produced with ESM2-650M.^11^The digestion-defining boundary residues were held fixed. In `esm2_p`*k, k*% of the mutable residues were masked and replaced with the highest-probability amino acid satisfying the generator constraints. In `esm2_n_c`, the leftmost and rightmost the mutable residues were mutated with the same constrained replacement rule. One-sided variants (`esm2_n` and `esm2_c`) were also evaluated.

All generators can be implemented either at the peptide level or at the protein level. In the former case, decoy peptides are supplied directly to the search engine; in the latter, mutated peptides are concatenated back into decoy proteins and provided as a second protein set. Throughout this work, we use protein-level decoys for all the generators except for `Sage` that is implemented on a peptide level.

Decoy generation algorithms are described in more detail in Supplementary Note S1.

### Sequence-only separability audit

For each decoy family, compact neural-network classifiers were trained to predict target versus decoy labels from peptide sequence alone. Performance was summarized with 5-fold cross-validated AUC. Architecture, feature encoding, data splits, and hyperparameters are provided in the Supplementary Note S2.

### Spectral-space diagnostics

Each peptide was represented by a Prosit-predicted spectrum. ^16^Each spectrum is associated with a precursor mass *m*(·). Similarity between spectra was measured using cosine distance, but only for pairs whose precursor masses differ by less than a tolerance *ε*. Specifically, for spectra *s*_1_ and *s*_2_ we defined

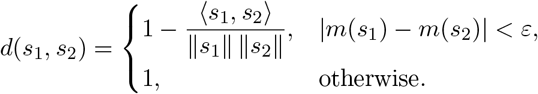

For a pepitde *p* we define Prosit-predicted spectrum as *ϕ*(*p*). Let *T* be the set of target peptides and *D* be the set of decoy peptides.

For a query spectrum *s*, the nearest-neighbor diagnostic was defined as

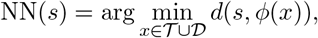

and the label of NN(*s*) was recorded. For a query peptide *p* it was defined as

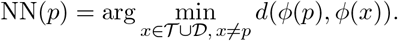

For target query *t* ∈ *T*, the nearest-decoy distance was defined as

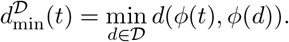

These diagnostics were analyzed overall and stratified by peptide length. Exact implementation details and plotting conventions are given in the Supplementary Note S3.

### End-to-end benchmarking

#### Pipelines

All end-to-end comparisons were run with the following analysis pipelines: Sage search ^15^and Sage followed by Oktoberfest rescoring. ^17^We selected these tools because both are open-source, which let us verify that our custom decoys were used correctly, and because they are easy to use and scale to the size of our study.

#### Reported metrics

At fixed q-value thresholds, we reported accepted identification counts and score distributions. In entrapment-enabled experiments, we additionally reported foreign-identification counts and false discovery proportion (FDP) lower and upper bounds following Wen *et al*.^18^With *N*_*T*_ accepted identifications from the original target search space, *N*_*E*_ accepted identifications from the foreign search space, *N*_*T* +*E*_ = *N*_*T*_ + *N*_*E*_, and effective foreign-to-original search-space ratio *r*, the bounds were

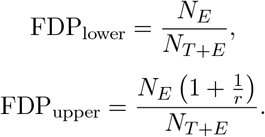

The evaluation pipeline is explained in more details in Supplementary Note S5.

## Results and Discussion

### Decoy adequacy requires global plausibility and local non-competitiveness

A useful way to think about decoy adequacy is that decoys are constrained by two competing design pressures. From a distance, the decoy database should look realistic enough that null spectra cannot systematically prefer targets over decoys (Figure 1B). Up close, however, a decoy should not imitate a particular target so well that it steals a true identification. These two ideas are simple to state, but it is helpful to write them down because they clarify why decoy design is fundamentally a balancing problem rather than an optimization in which “harder to distinguish is always better”.

For accepted PSMs at a threshold *k*, let *t* = *t*_*TP*_ + *t*_*FP*_ denote accepted target hits, where*t*_*TP*_ and *t*_*FP*_ are accepted true-positive and false-positive targets, and let *d* denote accepted decoy hits. Then

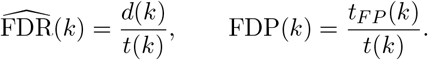

TDC is well calibrated only if decoy hits reproduce the behavior of false target hits (i.e. *t*_*FP*_ ≈ *d*). Rewriting this requirement in probabilistic terms highlights two competing requirements for practical decoy design.

#### Criterion 1: null exchangeability

For spectra *N* that do not correspond to any peptide in the target database, the chances of matching to target or decoy peptides are equal,

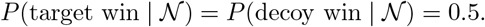

More precisely, this 0.5/0.5 symmetry is the idealized balanced case. In practice, it should be interpreted after accounting for the effective target-to-decoy library size ratio, since duplicate peptides, target–decoy overlaps, digestion constraints, and mass filtering can make the two candidate spaces unequal.

Intuitively, this means that for spectra whose best-scoring target interpretation is incorrect, for example because the spectrum is noisy, the true peptide is not present in the library, or the search engine produces artifacts, incorrect target wins and decoy wins should be approximately exchangeable.

#### Criterion 2: target protection

For spectra *T* generated by true peptides in the database,

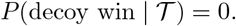

This is a safety condition. A decoy is allowed to be realistic, but it should not be so similar to a real peptide that it beats the correct target when the evidence actually comes from that target. A simple example of the opposite behavior is isobaric *I* ↔ *L* replacement. If the target peptide `PEPTIDE` is present in the library together with a decoy peptide `PEPTLDE`, the search engine will not be able to reliably distinguish between them.

In this form, the condition is idealized, since occasional matches are always possible. Instead of aiming to make this probability exactly zero, one should aim to reduce it as much as possible while not violating Criterion 1.

Together, these assumptions expose the central tension of decoy design. Criterion 1 asks decoys to resemble targets at the population level, whereas Criterion 2 asks decoys not to resemble any specific target too closely. Importantly, the corresponding probabilities are not properties of a decoy generator alone. They depend on the search engine, the data distribution, the target database, and the decoy database. Consequently, decoy generation is better treated as a fitting problem than as the search for a universally best generator. We therefore evaluate generators with a hierarchy of diagnostics that probe different aspects of decoy adequacy.

### Sequence-only models detect generator fingerprints

Initially, we ask a deliberately narrow question: can targets and decoys be distinguished from sequence alone, without seeing any spectrum at all? This does not directly measure search quality, but it provides a useful first-pass audit for generator artifacts. If a simple classifier can easily separate targets from decoys, then the generator leaves a detectable fingerprint in sequence space, which could potentially be exploited by a scoring model.

To address this question, we train a relatively simple binary classifier that takes peptide sequences as input (Figure S1A) and test its ability to separate targets from decoys generated by different methods (Figure 2). As a control, we first ask whether the classifier can separate two sets of target peptides. For this purpose, the human proteome is randomly split into two equal halves, and the two subsets are treated as target and decoy. As expected the resulting AUC of 0.5 indicates that the classifier does not show an obvious tendency to overfit and provides a reasonable baseline for the subsequent comparisons. We then evaluate decoys generated by reverse, shuffle, and mutation-based rules such as the DIA-NN-style generator (Table S2), as well as several ESM2-based strategies. For the ESM2 model (650M parameters), we test edits at the C-terminus, N-terminus, both termini (NC-terminus), and masking of different fractions of the peptide length (10%, 20%, and 30%). Most ESM2-based variants are harder to separate from targets than classical reverse decoys, although this advantage becomes weaker for the NC-terminus and 30% masking settings. In addition, the C-terminal ESM2 variant is substantially harder to distinguish than the DIA-NN decoys (AUC 0.64 versus 0.91), further highlighting the vulnerability of fixed substitution rules. The DIA-NN decoys are the easiest to separate even though they are purposely designed to introduce fragment-mass perturbations (Methods), with peptide sequences that the search engine never uses. This shows that construction-related signals can leak when peptide sequences or indirect hints are exposed without proper attention.

**Figure 2:**
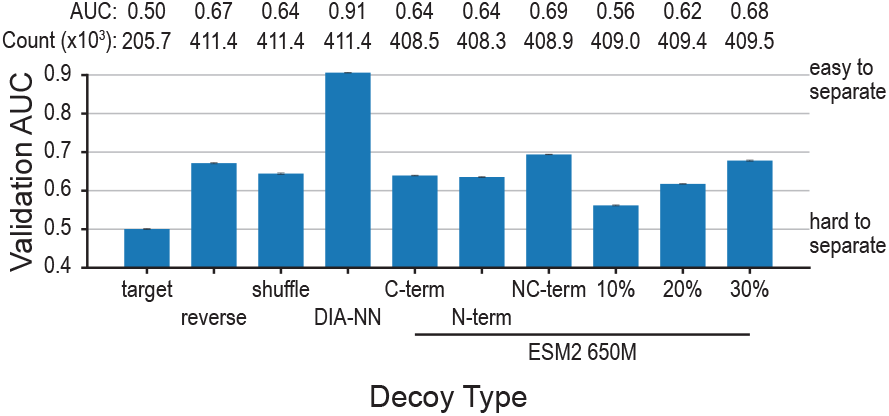
Target-decoy separability test. A sequence-only classifier was trained to distinguish target peptides from decoys generated by each method for the canonical human proteome. For the “target” control, the human proteome was split equally, and the two halves were treated as target and decoy. Bars show performance as area under the curve (AUC) on a held-out test set. Error bars indicate variation across 5-fold splits.

To place these results in context, we also test the classifier on peptides from other species, following the logic of entrapment-style benchmarks. Specifically, we compare human target peptides against peptides from *Mus musculus, Arabidopsis thaliana, Saccharomyces cerevisiae, Escherichia coli*, and Archaea (taxon ID 2157) (Figure S1B). As expected, peptides from evolutionarily more distant organisms are generally easier to separate from human peptides. The most extreme case was the Archaea, which reached an AUC of 0.83, likely reflecting differences in amino acid usage and sequence composition associated with adaptation to distinct environments. This experiment underscores that an entrapment proteome should not be chosen only by matching size. It should also be neither too similar to the target proteome, which would make separation unrealistically difficult, nor too distant, which would make it trivially easy.

Finally, because ESM2 is available in several model sizes (8M, 35M, 150M, 650M parameters), we ask whether model size affects target–decoy separability. Across repeated experiments spanning different model sizes and decoy-generation modes (Figure S1C), two consistent patterns emerge. First, decoys become easier to detect when more peptide positions are changed. Second, larger models generally produce decoys that are slightly harder to separate from targets across generation modes. This effect, however, is modest between 150M and 650M, whereas the computational cost of using larger models (e.g. 3B parameters) is substantially higher. We therefore use the 650M model for the downstream analyses.

Overall, the sequence-only experiments show that classical generators leave detectable fingerprints, whereas ESM2-based decoys are harder to distinguish from targets. This suggests that PLM-based generation reduces obvious sequence-level artifacts. At the same time, the practical meaning of this result should not be overstated. Sequence-only classification is a leakage screen, not a stand-alone criterion of decoy adequacy for proteomics analysis.

### Spectral-space diagnostics reveal balanced null behavior and short-peptide collisions

We next examine decoy behavior in a representation that is more closely tied to peptide distinguishability in mass spectrometry. Each peptide is represented by a Prosit-predicted spectrum, ^16^and proximity is measured by cosine distance. If two peptides are very close in this space, they are difficult to distinguish reliably during database search. Although this representation does not reproduce any specific search-engine score, it provides a useful search-engine-agnostic proxy for local peptide distinguishability.

We probe local spectral neighborhoods around three types of queries: spectra corresponding to target peptides, spectra corresponding to decoy peptides, and randomly generated spectra intended to approximate noise (Figure 3A, Supplementary Note S3). This allows us to examine how targets and decoys populate nearby spectral space and whether local competition remains balanced between the two classes. If “null exchangeability” holds approximately, nearby competitors should not systematically favor one class over the other. We therefore use nearest-neighbor labels and local target-win rates as intuitive proxies. The results are summarized in Figure 3B–C, with additional ESM2 settings shown in Figure S2. Notably, decoy families differ markedly in how well they preserve this balance. Reverse and shuffle decoys show a clear asymmetry: target queries tend to prefer reverse or shuffle decoys, whereas reverse or shuffle decoy queries tend to prefer targets. This reciprocal bias is inconsistent with “null exchangeability” and indicates that these generators distort local competitive structure in spectral space. In contrast, the DIA-NN-style generator and especially ESM2-based decoys remain much closer to the locally balanced regime. Thus, these generators better preserve local competitive structure in spectral space, which is the aspect most relevant to search and rescoring. Finally, when probing neighborhoods around randomly generated spectra intended to approximate noise, we observe that all decoy generators exhibit well-balanced local structure (Figure 3D, Supplementary Note S3). In this setting, nearest-neighbor labels and local target-win rates remain close to the expected proportions given the global target-to-decoy ratio, with no systematic preference for either class. This suggests that, in the absence of meaningful peptide signal, all generators behave consistently with approximate “null exchangeability”. The lack of detectable asymmetry for random queries further indicates that the imbalances observed for target and decoy queries arise from how different generators structure peptide-like spectral neighborhoods, rather than from global artifacts of the search space.

**Figure 3:**
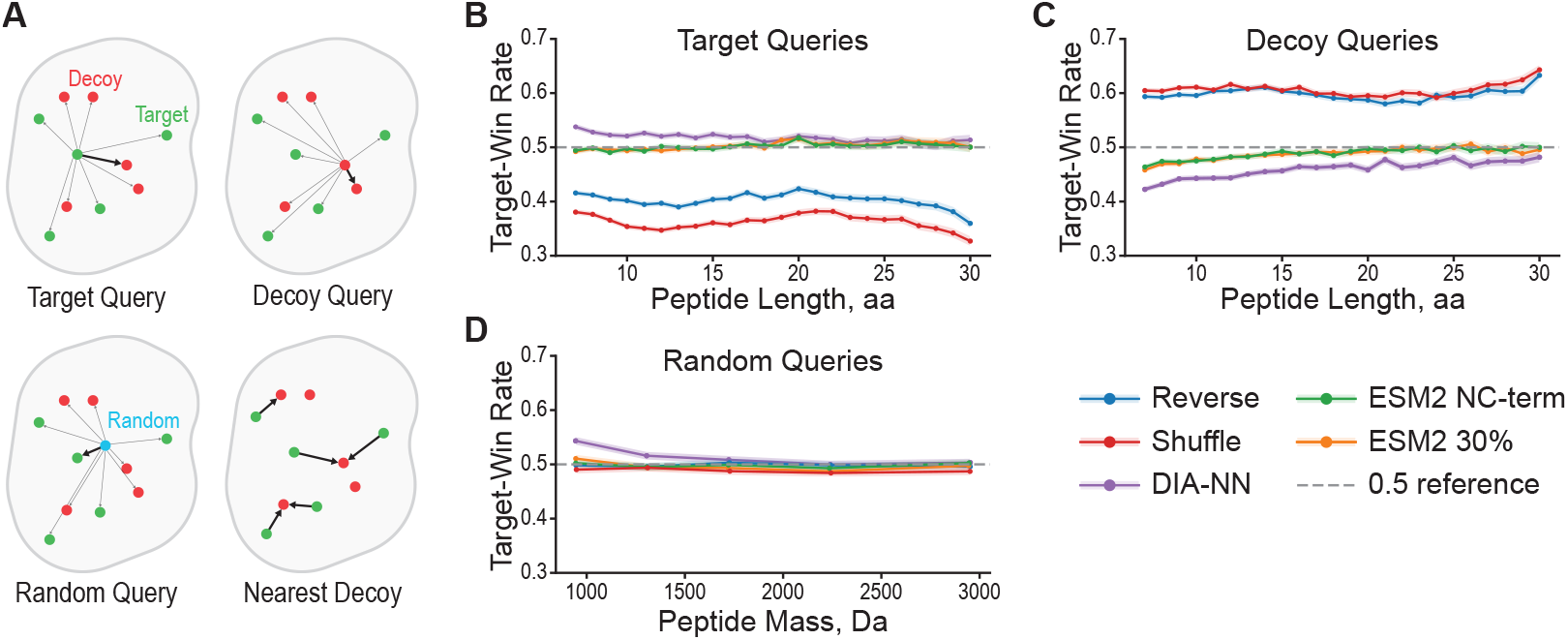
“Null competition” analysis of decoy generation strategies. Each panel summarizes the effective target-win rate, defined as the fraction of queries for which the best-scoring false-target competitor outscored the best-scoring decoy competitor by spectral cosine similarity. **(A)** Schematic overview of the query types used in the analysis. **(B)** Target-win rate as a function of query peptide length for target-peptide queries. **(C)** The same metric for decoy-peptide queries. **(D)** Target-win rate as a function of query precursor mass for random spectra. Five decoy generation strategies are compared: Reverse, Shuffle, DIA-NN, ESM2 30%, and ESM2 NC-term. Shaded bands denote 95% Wilson confidence intervals. The dashed line indicates a baseline (0.5) for a perfectly calibrated decoy generator.

A potential explanation for the observed target–decoy asymmetry in the reverse and shuffle generators is their isobaric construction: because these decoys share the precursor mass of their corresponding targets, one might expect the nearest neighbor of a spectrum from one class to often be its paired counterpart from the other class. This effect may indeed contribute to the observed behavior. Consistent with this idea, in our experiment the closest decoy to a given peptide was its matched decoy for about 5–10% of peptides for the reverse and shuffle generators, which is of the same order as the observed imbalance (Figure S5, Supplementary Note S4). At the same time, this mechanism alone may not provide a complete explanation. A similar fraction is also observed for the DIA-NN generator, but the asymmetry is not present there. Notably, DIA-NN decoys constructed on a peptide level are not isobaric. In our setting, however, decoys are constructed at the protein level, so isobaric target–decoy replacements can still arise indirectly through miscleaved peptides (Supplementary Note S1). Taken together, these observations suggest that direct target–decoy pairing may be part of the explanation, but that broader distortions in how reverse and shuffle decoys populate local spectral neighborhoods are also likely to play a role.

We next consider “target protection” by asking how close a target peptide lies to its nearest decoy neighbor. For this purpose, we use the nearest-decoy distance 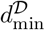 as an intuitive proxy. If a target has a decoy extremely close in spectral space, it resides in a collision-prone neighborhood, and the corresponding spectra may be vulnerable to matching the decoy instead of the target. This effect is already evident for the reverse generator, where short peptides often have highly similar reversed counterparts (Figure 4A). More broadly, the dominant structural pattern across all generators is peptide-length dependence, which is in line with earlier observations from database-search scoring: short peptides are intrinsically harder to separate from incorrect matches. ^19^As shown in Figure 4B–D, the distance between a target and its nearest decoy decreases markedly for shorter peptides, with the strongest crowding observed around lengths 7–9. At the same time, the severity of this effect differs across generators, indicating that decoy design influences how strongly short-peptide neighborhoods become collision-prone. A more detailed analysis for peptide lengths 7–12 across different generators is shown in Figure S3.

**Figure 4:**
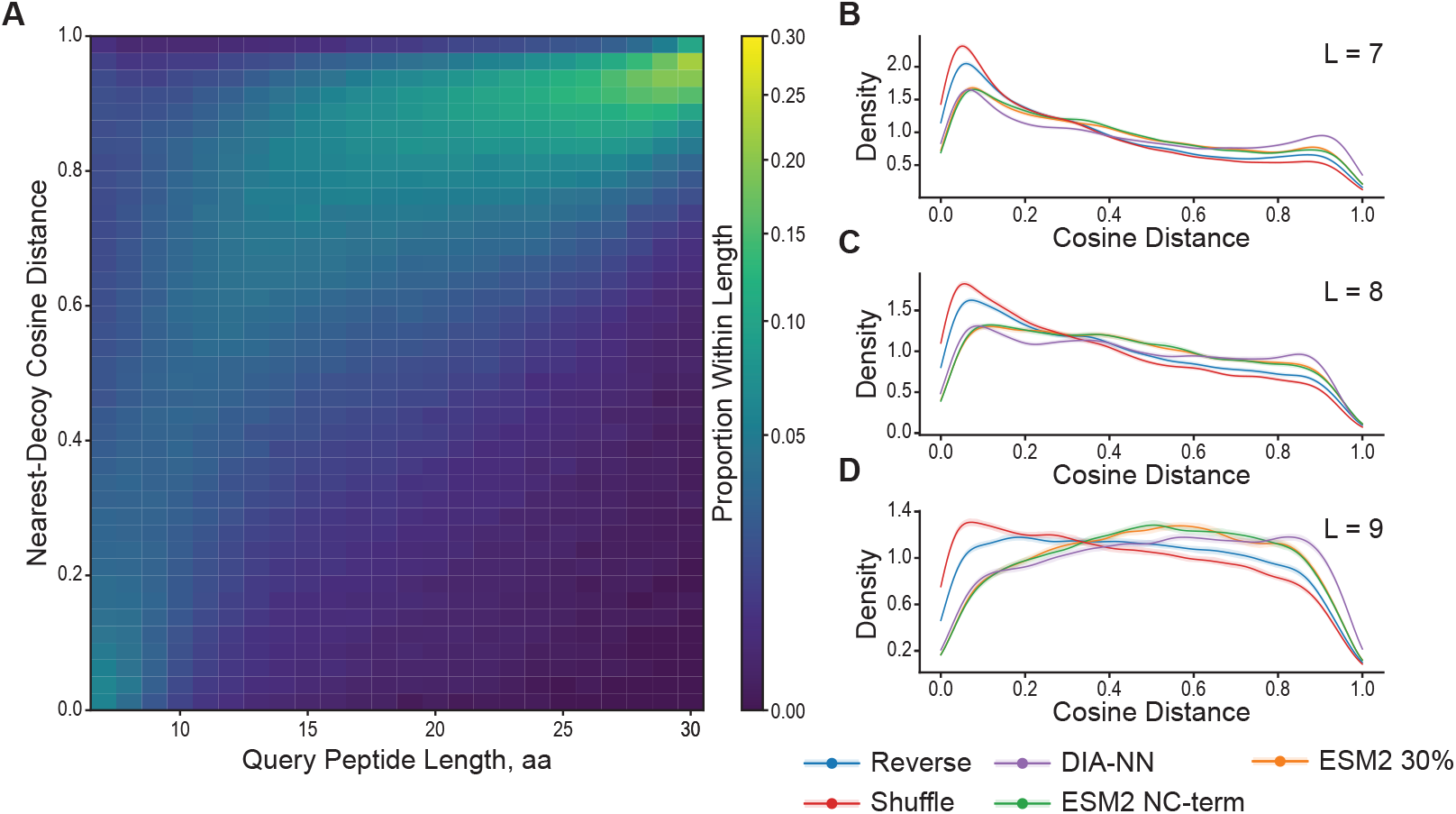
Nearest-decoy distance analysis of “target protection”. For every target peptide, the cosine distance to its closest decoy within the precursor mass window quantifies how dissimilar the best available decoy spectrum is; lower values indicate stealthier decoys that more closely mimic a true target match. **(A)** Heatmaps of the nearest-decoy cosine distance distribution conditioned on target peptide length for the reverse strategy. **(B-D)** Kernel density estimates of the nearest-decoy cosine distance distribution at peptide lengths 7, 8, and 9, respectively, overlaying Reverse, Shuffle, DIA-NN, ESM2 30%, and ESM2 NC-term decoy strategies. Shaded bands indicate 95% bootstrap confidence intervals.

This trend is important because aggregate summaries can easily obscure it. Short peptides occupy a much smaller combinatorial space, so as peptide libraries grow, close target–decoy neighbors become increasingly difficult to avoid. In this sense, short peptides define an intrinsically collision-prone regime rather than merely exposing an isolated weakness of one generator. Even among targets alone, sufficiently large databases inevitably contain pairs of peptides that are very close in sequence or spectral representation. A generator may therefore appear acceptable on average while still producing problematic local neighborhoods for short peptides. We therefore recommend reporting nearest-decoy diagnostics not only globally, but also as a function of peptide length.

Inspection of the closest reverse target–decoy pairs makes this geometric picture more concrete. Many of the nearest collisions are not exotic failures, but chemically or combinatorially simple transformations, especially *I* ↔ *L* equivalence and local *XY* ↔ *Y X* swaps (Figure S4B–G, Supplementary Note S4). This observation helps translate an abstract distance-based diagnostic into recognizable peptide-level motifs and explains why reverse decoys can remain a strong aggregate baseline while still producing vulnerable local neighborhoods for specific peptides.

Taken together, these spectral-space diagnostics provide a more mass-spectrometry-relevant view of decoy adequacy. They support two conclusions. First, ESM2-based generators preserve local target–decoy neighborhoods more faithfully than the other tested methods and therefore behave most favorably with respect to “null exchangeability”. Second, the strongest threat to “target protection” arises in the short-peptide regime, where crowded local structure makes close target–decoy collisions difficult to avoid for any realistic generator.

### Stress-test generators provide interpretable extremes

The synthetic `Random` and `Isobaric` generators were designed as diagnostic anchors rather than practical production choices. `Random` represents the “too easy” extreme: targets and decoys are trivially distinguishable, so any diagnostic should flag strong target/decoy separation. `Isobaric` represents the opposite extreme: decoys become so close to real targets that target protection begins to fail. These anchors are valuable because they make the intermediate behavior of reverse, shuffle, and ESM-based generators much easier to interpret. We evaluate these generators using a Sage search on yeast data augmented with maize entrapment, which provides an external proxy for false discoveries. This setup allows us to simultaneously inspect the shape of score distributions and compare nominal FDR with empirical FDP across generators (Figure 5).

**Figure 5:**
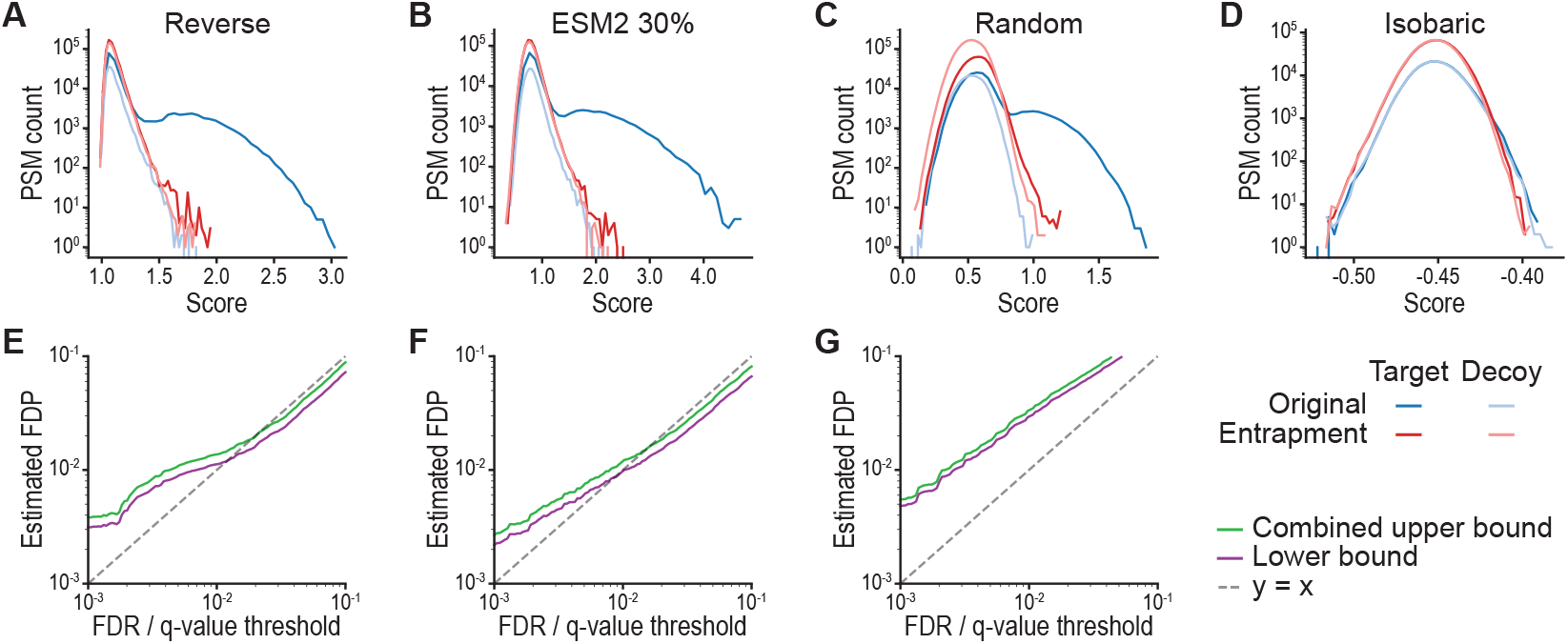
Score distributions and entrapment-based calibration across stress-test decoy generators. **(A–D)** PSM score distributions for Reverse, ESM2 650M 30%, Random, and Isobaric decoys. **(E–G)** Entrapment-based empirical FDP versus nominal FDR curves, with lower and upper bounds, ^18^for Reverse, ESM2 650M 30%, and Random. Isobaric generator is omitted since it produces no reliable identifications.

For `Reverse` decoys, we observe the characteristic bimodal distribution of target scores, with one mode corresponding to lower-scoring matches and a second mode corresponding to high-scoring matches. Decoy PSMs, as well as both target and decoy matches from the entrapment database, populate primarily the lower-scoring regime, yielding similar distributions across these groups (Figure 5A). ESM2-based decoys at 30% masking exhibit a closely related structure (Figure 5B), with a comparable separation between low- and high-scoring matches and similar alignment between primary and entrapment-derived distributions.

In contrast, `Random` decoys alter the shape of the score distribution (Figure 5C). While two modes are still present, they are less clearly separated, and the overall distribution is skewed relative to reverse or ESM2 decoys. Decoy PSMs are concentrated at lower scores, and the relative positioning of target and decoy distributions differs from that in the previous cases.

At the other extreme, `Isobaric` generators produce nearly indistinguishable score distributions across all classes (Figure 5D). Targets and decoys from both the original and entrapment databases follow the same overall distribution, with only a slight shift between primary and entrapment targets, reflecting differences in how well peptides from each source match the observed spectra.

These differences in score structure are reflected in the entrapment-based calibration curves (Figure 5E–G). Reverse and ESM2 decoys show close tracking between estimated FDR and empirical FDP across most of the range, with small deviations at low FDR thresholds. `Random` decoys, by contrast, exhibit larger discrepancies between these quantities, particularly at lower thresholds. For the `Isobaric` generator, no stable set of identifications is obtained, and the calibration curves are therefore not shown.

Together, these stress-test generators illustrate how systematic changes in decoy construction translate into predictable changes in both score distributions and entrapment-based calibration, providing interpretable reference points for evaluating more realistic decoy designs.

### End-to-end benchmarks show limited gains over reverse under current search settings

We next evaluate decoy generators in full search and rescoring pipelines, where their impact on identification rates and calibration can be assessed directly. We benchmark a range of generators, including reverse, shuffle, DIA-NN-style, and ESM2-based decoys, across multiple datasets and analysis settings using Sage with and without Oktoberfest rescoring (Figures 6, S6, S7, S8, Supplementary Note S6).

**Figure 6:**
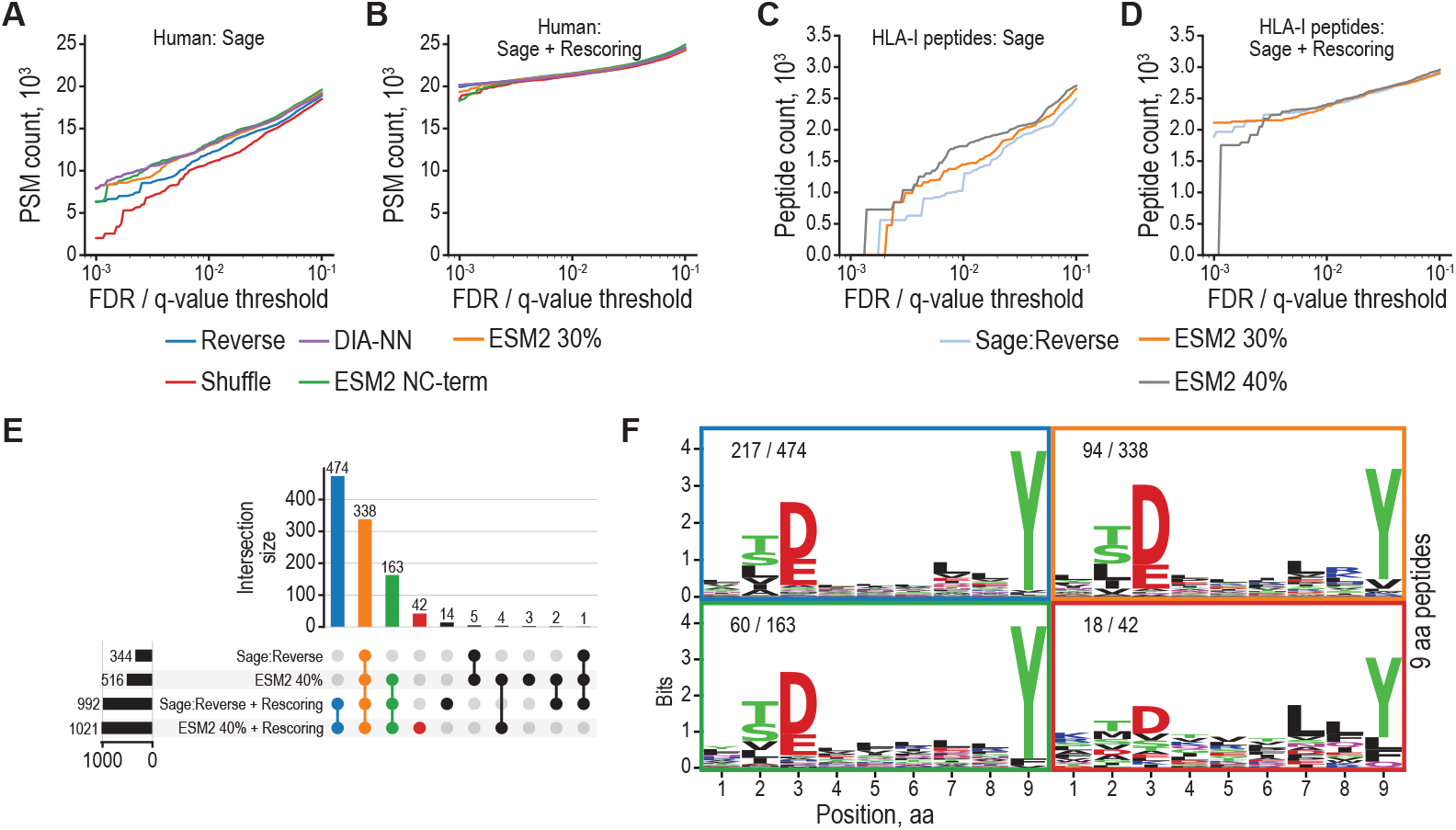
Impact of decoy generation and rescoring on proteome and immunopeptidome identifications. **(A–B)** Cumulative PSM counts versus q-value for the human cell line proteome^7^without **(A)** and with **(B)** Oktoberfest rescoring. ^17^**(C–D)** Cumulative peptide count versus q-value for HLA-A*01:01 immunopeptidomics ^20^without and with Oktoberfest rescoring. **(E)** Overlap of identified immunopeptides between Reverse and ESM2 650M 40% decoys, further stratified by rescoring option (as in **C–D**). **(F)** Sequence logo for 9-mer immunopeptides in the intersection sets shown in **(E)**, using the corresponding colors.

On a human proteome dataset analyzed with Sage, all generators except `DIA-NN` exhibit similar identification rates across a range of score thresholds (Figure 6A). DIA-NN-style decoys consistently yield fewer target identifications under the same conditions. After applying Oktoberfest rescoring, the total number of accepted identifications increases substantially for all generators, and the differences between `Reverse`, `Shuffle`, and ESM2-based decoys become even smaller (Figure 6B). `DIA-NN`-style decoys remain separated from the others in terms of identification rate.

We next consider a monoallelic HLA immunopeptidomics dataset (HLA-A*01:01),^20^where sequence constraints differ substantially from those in tryptic proteomics. In this setting, ESM2-based decoys at higher masking levels (40%) yield higher identification rates than both reverse decoys and lower masking levels when using Sage alone (Figure 6C). Following Oktoberfest rescoring, the total number of identified peptides increases across all generators, and the relative differences between them are reduced (Figure 6D). Entrapment-based analysis of HLA immunopeptidomics search is provided in Figure S9.

To further examine differences between generators, we compare peptide-level overlaps using an UpSet analysis (Figure 6E). Considering ESM2 (40%) and reverse decoys, with and without rescoring, most peptides uniquely identified by ESM2 in the non-rescored setting are also present in the union of identifications after rescoring. Finally, sequence logos for 9-mer peptides in the set difference between ESM2 and reverse decoys (without rescoring) show a consistent motif structure (Figure 6F).

Overall, while individual settings may show modest differences between generators, the aggregate behavior across datasets and pipelines indicates that improvements over reverse decoys are limited and context-dependent under current search configurations. Using post-search rescoring improves identification rates for all decoy generators and reduces the differences in performance between them.

## Conclusions

Here, we revisited decoy adequacy in target–decoy competition and introduced a PLM-based strategy for decoy generation. Across the settings tested here, PLM-based decoys did not consistently improve end-to-end identification performance over strong classical baselines such as reverse decoys. At the same time, they behaved favorably across several diagnostics, suggesting that their main value at present lies not in universal replacement of classical generators, but in providing tunable and target-like alternatives for benchmarking and stress testing.

A second contribution of this work is a practical framework for evaluating decoy strategies across three complementary settings – sequence-only target–decoy separability, search-engine-agnostic spectral-space diagnostics, and end-to-end search benchmarks with and without rescoring. Together, these analyses show that no single diagnostic is sufficient on its own. Sequence-only models mainly detect generator fingerprints, spectral-space analyses reveal local target–decoy geometry and collision-prone regimes, and full search pipelines determine whether these effects matter in practice. In particular, short peptides consistently emerge as the most vulnerable regime, with crowded local neighborhoods and frequent near-collisions.

These findings suggest that the choice of an appropriate decoy generator is inherently dependent on the data set, search space, and analysis pipeline. Rather than searching for a universally best generator, future work may benefit from optimizing decoy generation adaptively, for example by comparing multiple generators within the same workflow and selecting the one that is best suited to a given analysis setting (Supplementary Note S7). More broadly, we view decoy generation not only as a technical component of FDR estimation, but also as a diagnostic tool for understanding and stress-testing proteomics search engines as they become increasingly data-driven and increasingly reliant on machine-learning models.

## Abbreviations

TDC: target–decoy competition
FDR: false discovery rate
FDP: false discovery proportion
PSM: peptide-spectrum match
PLM: protein language model
ESM: Evolutionary Scale Modeling
ML: machine learning
DDA and DIA: data-dependent and data-independent acquisition.

## Data and Code Availability

**Repository:** https://github.com/SinitcynLab/DecoyGeneration. **Datasets:** accession identifiers and per-regime processing manifests are provided in the repository. The analyses in this study used the following data sets: *Saccharomyces cerevisiae* data set (ProteomeXchange: PXD044981), with optional maize entrapment;^21^human HepG2 cell line data set (ProteomeXchange: PXD022582); ^7^and mono-allelic human HLA peptide data set (MassIVE: MSV000080527). ^20^**Reproducibility:** exact software versions, configuration files, and command lines are included in the (Supplementary Notes).

## AI Use Disclosure

AI-assisted technologies were used in the preparation of this manuscript and in software development. ChatGPT (OpenAI) was used to assist with English grammar refinement. The conceptual development, scientific interpretation, and conclusions remain the sole responsibility of the authors. In addition, parts of the research code were generated with the assistance of ChatGPT (OpenAI) and Claude Code (Anthropic). All AI-generated code was carefully reviewed, tested, and validated by the authors before inclusion in the research workflow.

## Acknowledgment

We thank Jürgen Cox and Peter Cimermančič for discussions that helped shape the initial idea for this project and for reading the draft manuscript. We thank members of the AIT4Life and BioMS groups in Utrecht University for valuable feedback and discussions, in particular Douwe Schulte, Bas van Breukelen, Sanne Abeln and Albert J.R. Heck for reading the draft manuscript. We also acknowledge the “Trustworthiness in Proteomics” workshop at the Lorentz Center (Leiden, the Netherlands) for providing an important forum for scientific exchange, including discussions with Maike Weber, Mathias Wilhelm, Robbe Devreese, and Lukas Käll.

## Contribution

**Conceptualization:** G.R., M.M., P.S.; **Data curation:** G.R., F.K.; **Formal analysis:** G.R., M.M.; **Funding acquisition:** P.S.; **Investigation:** G.R., F.K.; **Methodology:** G.R., F.K., M.M., P.S.; **Project administration:** P.S.; **Resources:** H.W.P.v.d.T.; **Software:** G.R., F.K., M.M., P.S.; **Supervision:** M.M., H.W.P.v.d.T., P.S.; **Validation:** G.R., P.S.; **Visualization:** G.R., F.K., P.S.; **Writing – original draft:** G.R., F.K., M.M., P.S.; **Writing – review & editing:** all authors.

## Abstract Graphic

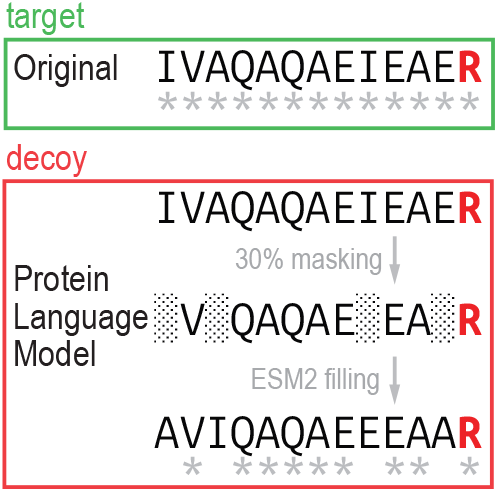

## References

[1] Elias, J. E.; Gygi, S. P. Target-decoy search strategy for increased confidence in large-scale protein identifications by mass spectrometry. Nature methods 2007, 4, 207–214.

[2] Elias, J. E.; Gygi, S. P. Target-decoy search strategy for mass spectrometry-based proteomics. 2009, 55–71.

[3] Burger, T. Gentle introduction to the statistical foundations of false discovery rate in quantitative proteomics. Journal of proteome research 2018, 17, 12–22.

[4] Käll, L.; Canterbury, J. D.; Weston, J.; Noble, W. S.; MacCoss, M. J. Semi-supervised learning for peptide identification from shotgun proteomics datasets. Nature methods 2007, 4, 923–925.

[5] Fondrie, W. E.; Noble, W. S. mokapot: Fast and flexible semisupervised learning for peptide detection. Journal of Proteome Research 2021, 20, 1966–1971.

[6] Demichev, V.; Messner, C. B.; Vernardis, S. I.; Lilley, K. S.; Ralser, M. DIA-NN: neural networks and interference correction enable deep proteome coverage in high throughput. Nature methods 2020, 17, 41–44.

[7] Sinitcyn, P.; Hamzeiy, H.; Salinas Soto, F.; Itzhak, D.; McCarthy, F.; Wichmann, C.; Steger, M.; Ohmayer, U.; Distler, U.; Kaspar-Schoenefeld, S.; others MaxDIA enables library-based and library-free data-independent acquisition proteomics. Nature biotechnology 2021, 39, 1563–1573.

[8] Adams, C.; Laukens, K.; Bittremieux, W.; Boonen, K. Machine learning-based peptide-spectrum match rescoring opens up the immunopeptidome. Proteomics 2024, 24, 2300336.

[9] Granholm, V.; Noble, W. S.; Käll, L. A cross-validation scheme for machine learning algorithms in shotgun proteomics. BMC bioinformatics 2012, 13, S3.

[10] Danilova, Y.; Voronkova, A.; Sulimov, P.; Kertész-Farkas, A. Bias in false discovery rate estimation in mass-spectrometry-based peptide identification. Journal of proteome research 2019, 18, 2354–2358.

[11] Lin, Z.; Akin, H.; Rao, R.; Hie, B.; Zhu, Z.; Lu, W.; Smetanin, N.; Verkuil, R.; Kabeli, O.; Shmueli, Y.; others Evolutionary-scale prediction of atomic-level protein structure with a language model. Science 2023, 379, 1123–1130.

[12] van Eck, J.; Gogishvili, D.; Silva, W.; Abeln, S. PLM-eXplain: divide and conquer the protein embedding space. Bioinformatics 2026, 42, btaf631.

[13] He, K.; Fu, Y.; Zeng, W.-F.; Luo, L.; Chi, H.; Liu, C.; Qing, L.-Y.; Sun, R.-X.; He, S.-M. A theoretical foundation of the target-decoy search strategy for false discovery rate control in proteomics. arXiv preprint arXiv:1501.00537 2015,

[14] Moosa, J. M.; Guan, S.; Moran, M. F.; Ma, B. Repeat-preserving decoy database for false discovery rate estimation in peptide identification. Journal of proteome research 2020, 19, 1029–1036.

[15] Lazear, M. R. Sage: an open-source tool for fast proteomics searching and quantification at scale. Journal of Proteome Research 2023, 22, 3652–3659.

[16] Gessulat, S.; Schmidt, T.; Zolg, D. P.; Samaras, P.; Schnatbaum, K.; Zerweck, J.; Knaute, T.; Rechenberger, J.; Delanghe, B.; Huhmer, A.; others Prosit: proteome-wide prediction of peptide tandem mass spectra by deep learning. Nature methods 2019, 16, 509–518.

[17] Picciani, M.; Gabriel, W.; Giurcoiu, V.-G.; Shouman, O.; Hamood, F.; Lautenbacher, L.; Jensen, C. B.; Müller, J.; Kalhor, M.; Soleymaniniya, A.; others Oktoberfest: Open-source spectral library generation and rescoring pipeline based on Prosit. Proteomics 2024, 24, 2300112.

[18] Wen, B.; Freestone, J.; Riffle, M.; MacCoss, M. J.; Noble, W. S.; Keich, U. Assessment of false discovery rate control in tandem mass spectrometry analysis using entrapment. Nature Methods 2025, 22, 1454–1463.

[19] Cox, J.; Mann, M. MaxQuant enables high peptide identification rates, individualized ppb-range mass accuracies and proteome-wide protein quantification. Nature biotechnology 2008, 26, 1367–1372.

[20] Sarkizova, S.; Klaeger, S.; Le, P. M.; Li, L. W.; Oliveira, G.; Keshishian, H.; Hartigan, C. R.; Zhang, W.; Braun, D. A.; Ligon, K. L.; others A large peptidome dataset improves HLA class I epitope prediction across most of the human population. Nature biotechnology 2020, 38, 199–209.

[21] Staes, A.; Mendes Maia, T.; Dufour, S.; Bouwmeester, R.; Gabriels, R.; Martens, L.; Gevaert, K.; Impens, F.; Devos, S. Benefit of in silico predicted spectral libraries in data-independent acquisition data analysis workflows. Journal of Proteome Research 2024, 23, 2078–2089.

