## Supplementary Material for "Protein Language Model Decoys for Target Decoy Competition in Proteomics: Quality Assessment and Benchmarks"

**Supplementary Figures** **2**
Figure S1. Target-Decoy Separability Test. 2
Figure S2. "Null competition" analysis of decoy generation strategies. 3
Figure S3. Nearest-decoy distance analysis of "target protection". 4
Figure S4. Inspection of closest target–decoy pairs. 5
Figure S5. Inspection of closest decoy type for every structure. 6
Figure S6. Entrapment-based calibration across decoy generators for yeast data. 7
Figure S7. Entrapment-based calibration across decoy generators for human data. 8
Figure S8. Comparison of search results with and without rescoring. 9
Figure S9. Entrapment-based calibration across decoy generators for human HLA data. 10

**Supplementary Tables** **11**
Table S1. Overview of decoy generators used in this study. 11
Table S2. Fixed residue substitutions used in the DIA-NN-style decoy generator. 11

**Supplementary Notes** **12**
Note S1. Decoy generators 12
Note S2. Sequence-based separation 13
Note S3. Spectral-space neighborhoods and random null spectra 14
Note S4. Closest target–decoy pairs classification 16
Note S5. End-to-end search pipeline 16
Note S6. End-to-end search benchmarks 18
Note S7. Limitations and next steps 18

**Supplementary References** **20**

### Supplementary Figures

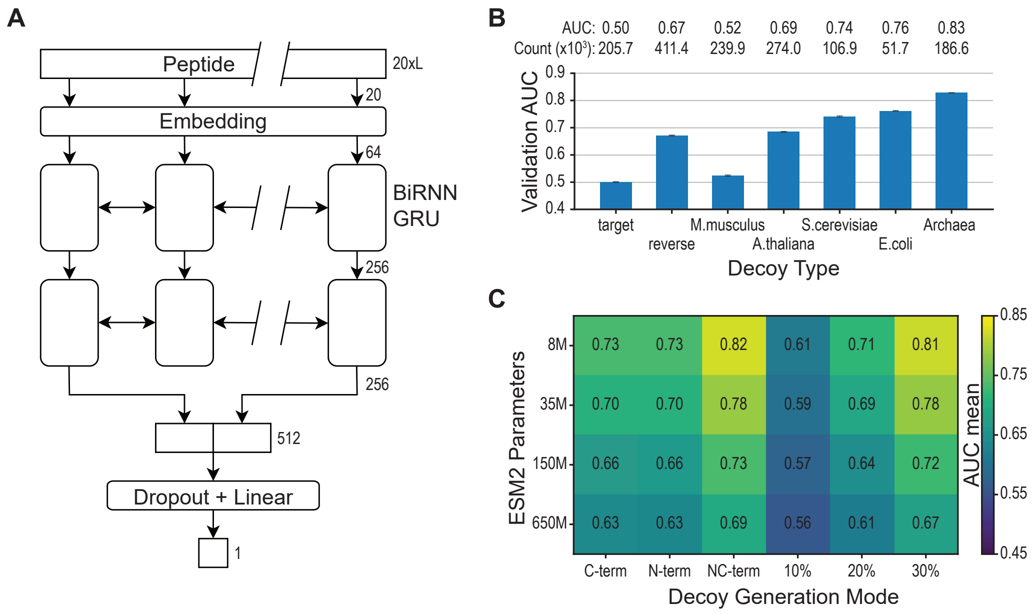


Figure S1: **Target-Decoy Separability Test.** **(A)** Schematic overview of the sequence-only target–decoy classifier. A detailed description of the model is provided in Supplementary Note S2. **(B)** Performance of sequence-only separation between the *Homo sapiens* proteome (Taxon ID: 9606) and proteomes from other species, including *Mus musculus* (Taxon ID: 10090), *Arabidopsis thaliana* (Taxon ID: 3702), *Saccharomyces cerevisiae* (Taxon ID: 559292), *Escherichia coli* (Taxon ID: 83333), and Archaea (Taxon ID: 2157). The results for the target and reverse controls are the same as in Figure 1 and are shown for a visual reference. **(C)** Performance of the classification model for different ESM2-based generation modes (C-term, N-term, NC-term, 10%, 20%, and 30%) across different ESM2 model sizes (8M, 35M, 150M, and 650M parameters).

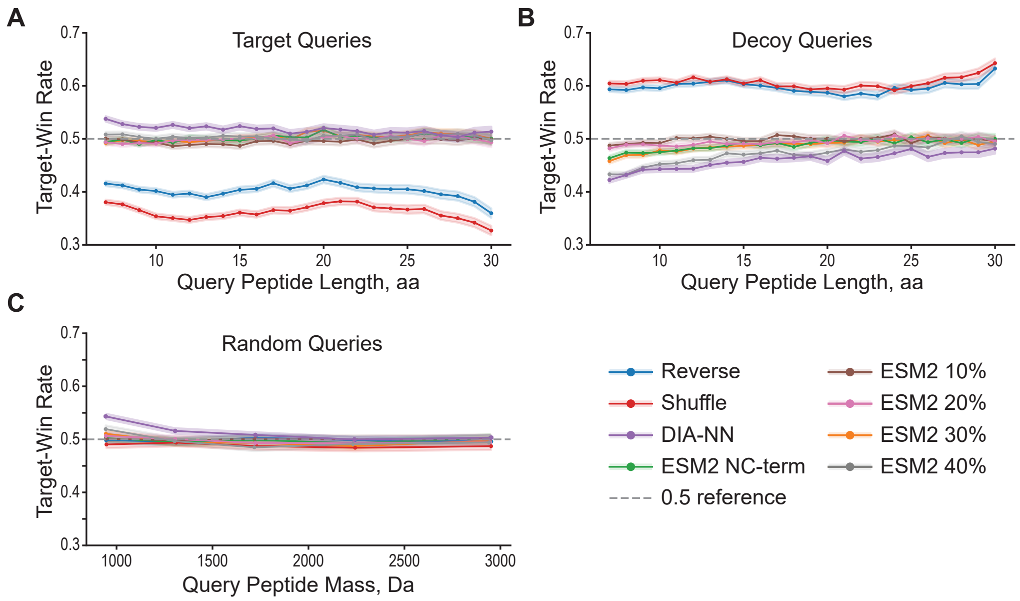


Figure S2: **"Null competition" analysis of decoy generation strategies.** Each panel summarizes the effective target-win rate, defined as the fraction of queries for which the best-scoring false-target competitor outscored the best-scoring decoy competitor by spectral cosine similarity. **(B)** Target-win rate as a function of query peptide length for target-peptide queries. **(C)** The same metric for decoy-peptide queries. **(D)** Target-win rate as a function of query precursor mass for random spectra. Eight decoy generation strategies are compared: Reverse, Shuffle, DIA-NN, ESM2 NC-term and ESM2 k% for $k = 10, 20, 30, 40, 50$. Shaded bands denote 95% Wilson confidence intervals. The dashed line indicates a baseline (0.5) for a perfectly calibrated decoy generator.

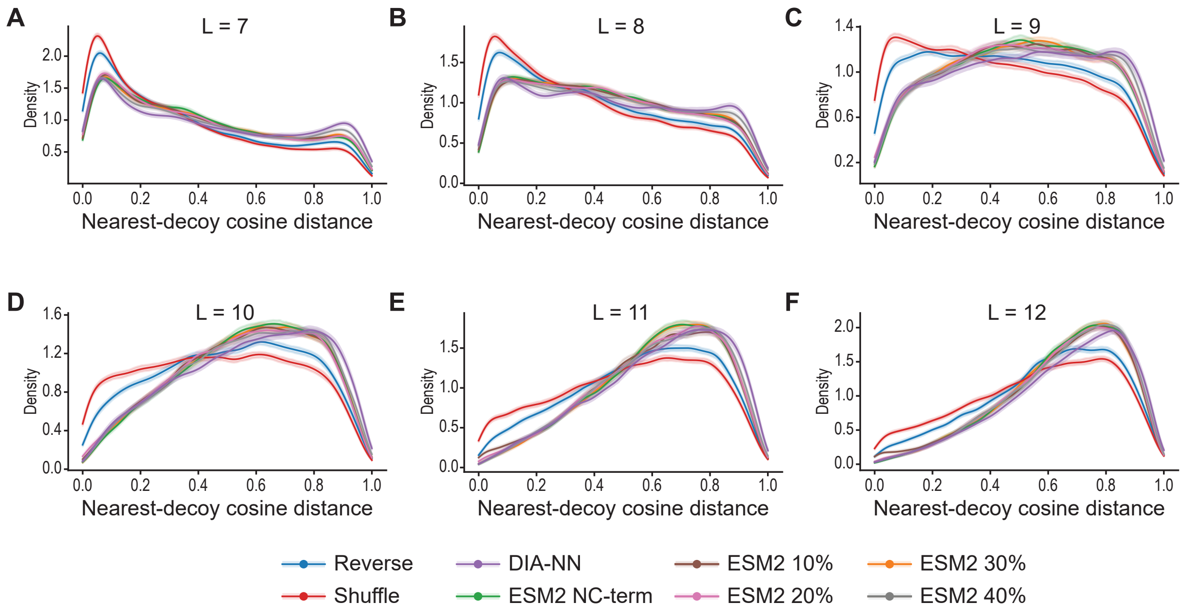


Figure S3: **Nearest-decoy distance analysis of "target protection".** For every target peptide, the cosine distance to its closest decoy within the precursor mass window quantifies how dissimilar the best available decoy spectrum is; lower values indicate stealthier decoys that more closely mimic a true target match. **(A-F)** Kernel density estimates of the nearest-decoy cosine distance distribution at peptide lengths 7–12, overlaying Reverse, Shuffle, DIA-NN, ESM2 NC-term, and ESM2 k% for $k = 10, 20, 30, 40, 50$ decoy strategies. Shaded bands indicate 95% bootstrap confidence intervals.

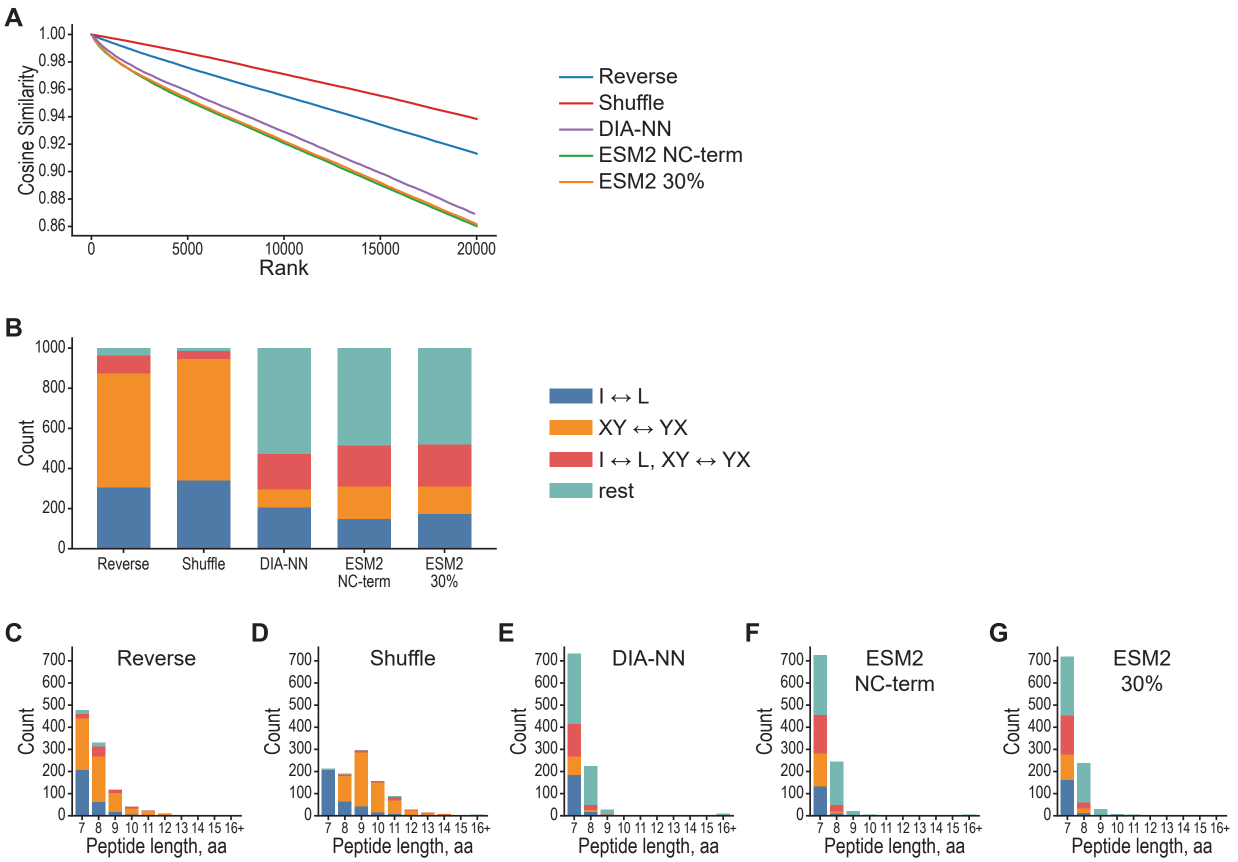


Figure S4: **Inspection of closest target–decoy pairs.** **(A)** Top 20,000 suspicious target–decoy pairs, visualized by ranking in ascending order of best cosine similarity. **(B)** Classification of the top 1,000 closest target–decoy pairs. **(C–G)** The same classification as in **(B)**, shown separately by peptide length.

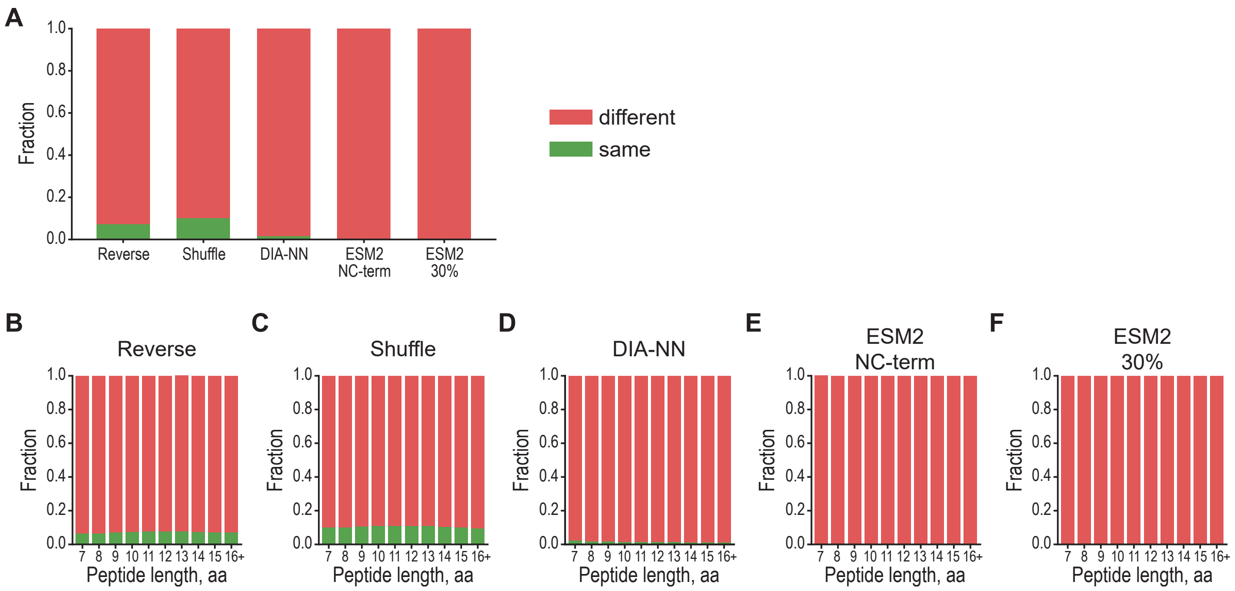


Figure S5: **Inspection of closest decoy type for every structure.** For every target peptide the closest decoy is identified and compared with decoy generated for the target. **(A)** Total fraction of target peptides whose closest decoy matches its own decoy across Reverse, Shuffle, DIA-NN, ESM2 NC-term and ESM2 30% generators. **(B)** The same classification as in **(A)**, shown separately by peptide length.

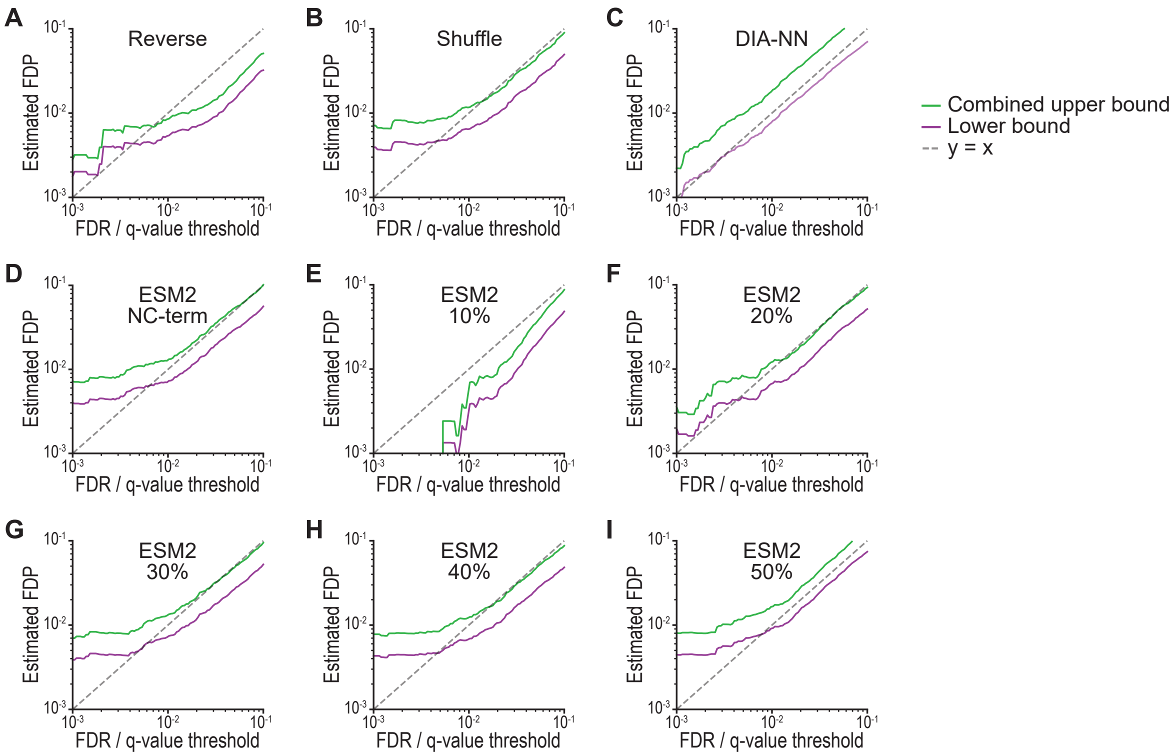


Figure S6: **Entrapment-based calibration across decoy generators for yeast data. (A–I)** Entrapment-based empirical FDP versus nominal FDR curves, with lower and upper bounds, for Reverse, Shuffle, DIA-NN, ESM2 NC-term and ESM2 k% for $k = 10, 20, 30, 40, 50$. Maize proteome is used as entrapment when searching over yeast data.

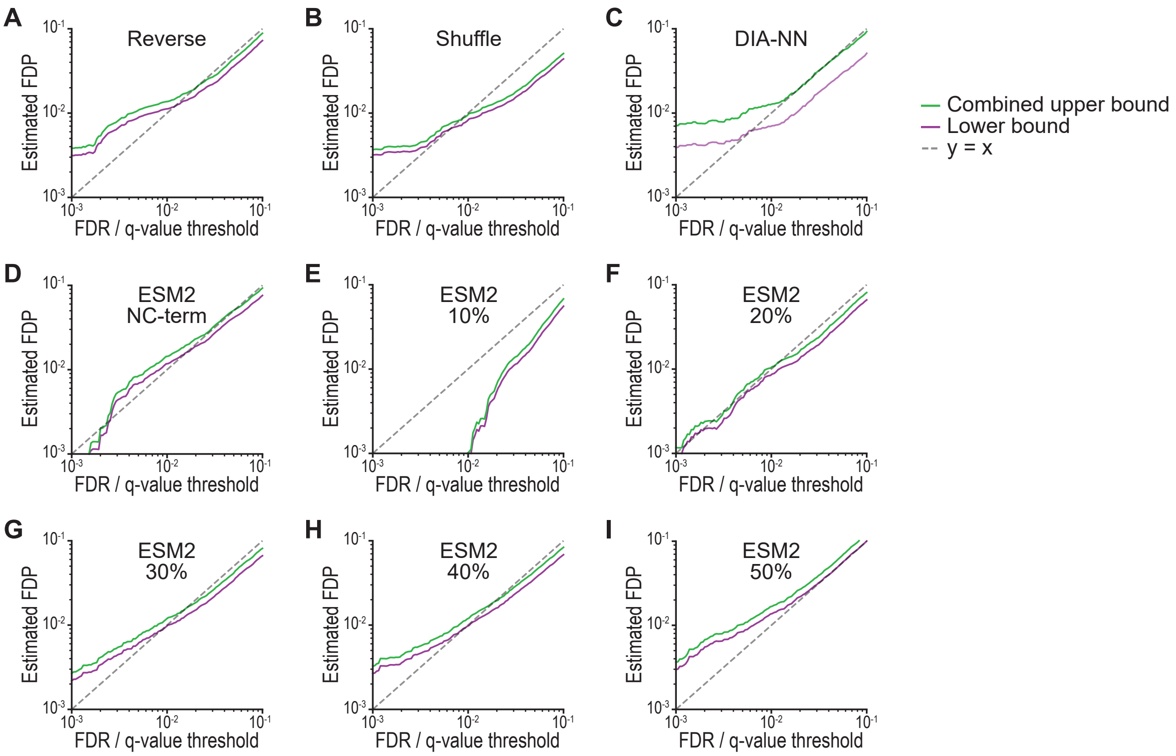


Figure S7: **Entrapment-based calibration across decoy generators for human data. (A–I)** Entrapment-based empirical FDP versus nominal FDR curves, with lower and upper bounds, for Reverse, Shuffle, DIA-NN, ESM2 NC-term and ESM2 k% for $k = 10, 20, 30, 40, 50$. Maize proteome is used as entrapment when searching over human data.

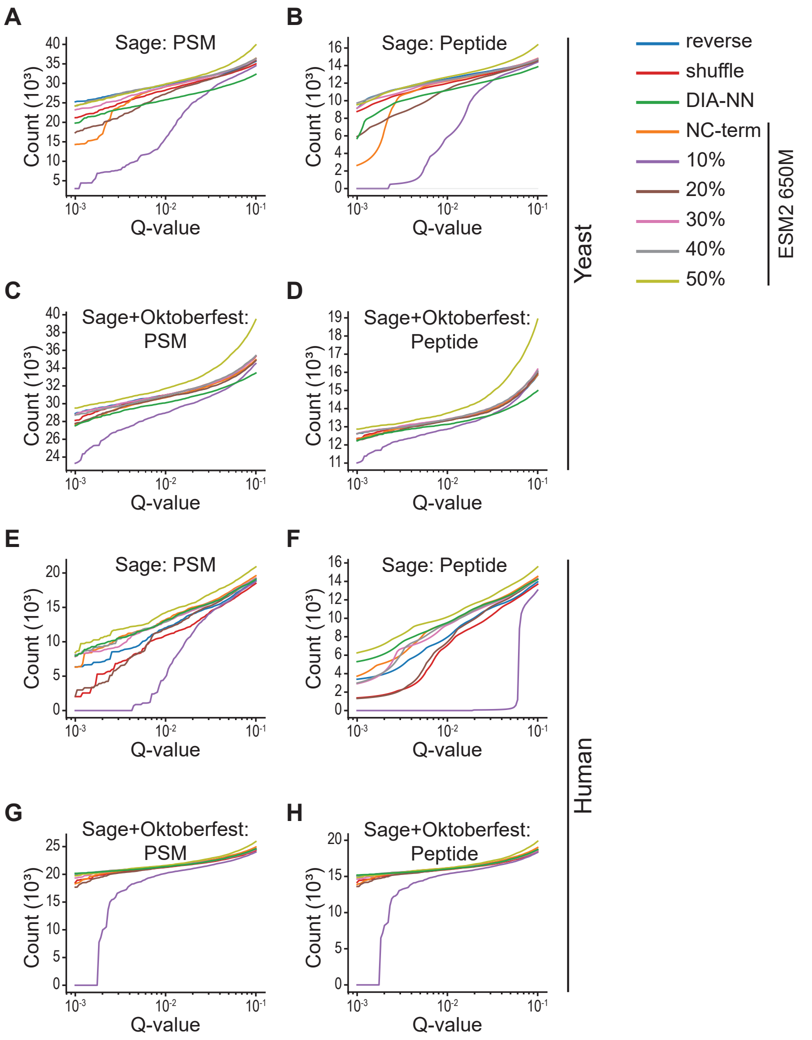


Figure S8: **Comparison of search results with and without rescoring.** **(A)** Yeast PSM-level results without rescoring. **(B)** Yeast peptide-level results without rescoring. **(C)** Yeast PSM-level results with Oktoberfest rescoring. **(D)** Yeast peptide-level results with Oktoberfest rescoring. **(E)** Human PSM-level results without rescoring. **(F)** Human peptide-level results without rescoring. **(G)** Human PSM-level results with Oktoberfest rescoring. **(H)** Human peptide-level results with Oktoberfest rescoring.

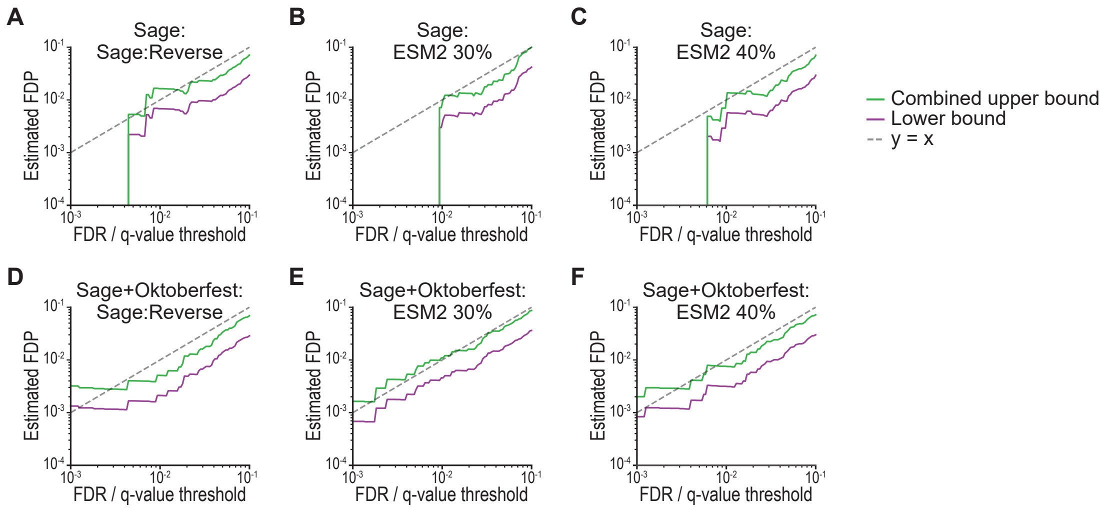


Figure S9: **Entrapment-based calibration across decoy generators for human HLA data. (A–C)** Entrapment-based empirical FDP versus nominal FDR curves, with lower and upper bounds, for Sage Reverse, ESM2 30%, and ESM 40% generators without rescoring. **(D–F)** Entrapment-based empirical FDP versus nominal FDR curves, with lower and upper bounds, for Sage Reverse, ESM2 30%, and ESM 40% generators with Oktoberfest rescoring. Maize proteome is used as entrapment when searching over human data.

### Supplementary Tables

Table S1: **Overview of decoy generators used in this study.** Examples assume a tryptic peptide, so the C-terminal cleavage-defining residue is kept fixed whenever the generator is digestion-aware. Illustrative outputs are schematic and are shown only to clarify the type of edit.

| Generator | Fixed positions | Transformation rule | Example / note |
|---|---|---|---|
| Reverse | Digestion-defining positions only | Reverse the mutable subsequence and write it back into the original mutable slots. | `PEPTIDER` → `EDITPEPR`. |
| Sage | First and last residue | Reverse the internal residues only. | `PEPTIDER` → `PEDITPER`. |
| Shuffle | Digestion-defining positions only | Randomly permute the mutable residues; identity permutations are rejected when possible. | `PEPTIDER` → `DPEIETPR`. |
| DIA-NN style | Digestion-defining positions only | Apply a local rule-based substitution at two positions, the first residue (index 0) and the residue immediately before the cleavage site, while preserving the digestion pattern. | Uses a fixed residue-substitution map $\rho(\cdot)$ defined in Table S2. |
| ESM-2 | Digestion-defining positions only | Mask selected mutable positions and replace each with the highest-probability amino acid proposed by ESM-2 that satisfies the generator constraints. | Actual output depends on model logits and the masked positions. |
| Random | None | Replace every peptide with an unrelated random peptide of length 15. | Deliberately unrealistic negative control. |
| Isobaric | Prefer edits away from digestion-defining positions | Apply near-isobaric edits with priority `I`↔`L`, then `GG`↔`N`, otherwise swap the first mutable adjacent pair. | Designed to create almost indistinguishable spectral neighbors. |

Table S2: **Fixed residue substitutions used in the DIA-NN-style decoy generator.**

| | | | | | | | | | | |
|---|---|---|---|---|---|---|---|---|---|---|
| G→L | A→L | V→L | L→V | I→V | F→L | M→L | P→L | W→L | S→T | C→S |
| T→S | Y→S | H→S | K→L | R→L | Q→N | E→D | N→Q | D→E | U→S | |

### Supplementary Notes

#### Note S1. Decoy generators

Let a peptide be denoted by $p = p_1 p_2 \dots p_n$. For digestion-aware generators, we distinguish a set of fixed positions $B \subseteq \{1, \dots, n\}$ and a complementary set of mutable positions $M = \{1, \dots, n\} \setminus B$. For a standard tryptic peptide, $B$ typically contains the C-terminal cleavage-defining residue when that residue is `K` or `R`. The generator is then applied only to the mutable subsequence while all fixed positions are copied back unchanged. This notation makes it possible to describe reverse, shuffle, and ESM-based generators in a uniform way.

The classical generators used here are lightweight sequence transformations. Reverse reverses the mutable subsequence while preserving the digestion-dependent positions. Sage follows the same general idea but additionally preserves the first residue, so only the internal part of the peptide is reversed. Shuffle replaces reversal by a random permutation of the mutable residues; in practice, identity permutations should be rejected whenever the peptide contains at least two mutable positions.

The DIA-NN-style generator is a local mutation heuristic rather than a global rearrangement. In words, it changes the residue adjacent to the digestion-specific boundary according to a fixed substitution rule while preserving the cleavage pattern. This produces decoys that remain peptide-like but are less obviously derived from full reversal or shuffling. Because the exact residue map was not included in the provided materials, we leave the final substitution table as an explicit placeholder to be filled with the rule used in the implementation.

The ESM-2 generator uses a protein language model to propose context-aware residue replacements. Mutable positions are masked, the model is queried on the masked peptide, and the highest-probability replacement satisfying the generator constraints is chosen. The constraints used in the manuscript are: (i) the replacement must differ from the original residue; (ii) `I` $\leftrightarrow$ `L` substitutions are disallowed; and (iii) the digestion pattern must not change. In the `esm2_pk` variant, $k\%$ of mutable positions are masked. In `esm2_n_c`, only the leftmost and rightmost mutable positions are edited. One-sided variants (`esm2_n` and `esm2_c`) edit only the N-side or only the C-side mutable position.

Two intentionally extreme generators were included as diagnostic anchors. Random replaces each peptide by an unrelated random 15-mer and therefore represents a deliberately easy negative class. Isobaric does the opposite: it tries to remain as close as possible to the target peptide. The procedure first attempts to apply `I`$\leftrightarrow$`L`. If that substitution is not applied, it then attempts to apply `GG`$\leftrightarrow$`N`. If neither substitution is applied, it performs the smallest available local swap of adjacent mutable residues. This produces decoys that are often spectrally almost indistinguishable from their targets and therefore acts as a stress test for target protection.

**ESM-2 constrained mutation.** The constrained replacement rule used in the ESM-2 family can be summarized as follows.

```
function ESM2ConstrainedDecoy(protein p, mask_positions Mmask):
    q <- p
    for position i in Mmask:
        logits <- ESM2(mask position i in q)
        for aa in amino_acids sorted by descending probability:
            if aa == p[i]:
                continue
            if {aa, p[i]} == {"I", "L"}:
                continue
```

```
            if changes_digestion_pattern(q, i, aa):
                continue
            q[i] <- aa
            break
    return q
```

**Isobaric stress-test generator.** The isobaric generator follows a fixed priority order designed to keep precursor mass and fragment patterns as similar as possible.

```
function IsobaricDecoy(peptide p):
    if p contains I or L at a mutable position:
        flip the first mutable I to L, otherwise the first mutable L to I
        return modified peptide

    if p contains GG at a mutable position pair:
        replace the first mutable GG by N
        return modified peptide

    if p contains N at a mutable position:
        replace the first mutable N by GG
        return modified peptide

    swap the first adjacent mutable pair that changes the sequence
    return modified peptide
```

All generators described here can be implemented either at the peptide level or at the protein level. In the peptide-level approach, in silico digestion (including potential missed cleavages) is performed prior to decoy generation; decoys are then generated independently for each peptide, and the resulting peptide library is used for database searching. In the protein-level approach, target proteins are first digested, the resulting peptides are mutated, and subsequently reassembled into protein sequences.

#### Note S2. Sequence-based separation

The sequence-based diagnostics uses a binary classifier that receives a peptide sequence as input and predicts whether it belongs to the target or the decoy set. The aim of this analysis is not to emulate a search engine, but to detect generator-specific fingerprints that are visible directly from sequence alone. Performance is summarized by cross-validated AUC.

**Input representation.** Peptides are treated as variable-length sequences over a fixed vocabulary consisting of the 20 standard amino acids together with the additional symbols `-BJOUXZ`. Two special tokens are added for padding and unknown characters. Each peptide is encoded as a sequence of token IDs and padded to the maximum sequence length within a minibatch.

**Model architecture.** The classifier is a bidirectional recurrent neural network (Figure S1A). In the manuscript runs, we use an embedding layer of dimension 64, followed by a two-layer bidirectional GRU with hidden size 256 per direction and dropout 0.2. The final forward and backward hidden states are concatenated and passed through a linear layer to produce a single logit, which is transformed by a sigmoid into a score between 0 and 1. In this convention, 1 corresponds to target and 0 to decoy.

**Training setup.** For each experiment, the target and decoy classes are first balanced by subsampling to the smaller class size. Exact peptide overlaps between target and decoy sets are removed before training. In the target-versus-target control, the human peptide set is shuffled and split into two equal halves, which are then treated as target and decoy. The model is trained with Adam, binary cross-entropy with logits, batch size 1024, learning rate $10^{-3}$, and 3 epochs.

**Evaluation.** Performance is estimated by 5-fold cross-validation. For each fold, the model is trained on four partitions and evaluated on the held-out partition. The manuscript reports the mean and standard deviation of validation AUC across folds. This setup is intentionally simple – the goal is to reveal sequence-level separability induced by decoy construction rather than to optimize predictive performance.

#### Note S3. Spectral-space neighborhoods and random null spectra

**Effective target and decoy peptide sets.** The workflow begins by digesting a target-decoy FASTA into unique peptides. Exact target/decoy sequence overlaps are not treated as genuine competitors: if the same peptide sequence appears in both target and decoy digests, it is promoted to the effective target set and removed from the effective decoy pool for competition analyses. This mirrors the practical convention that an exact target/decoy sequence tie should count as target rather than as a meaningful decoy alternative.

After digestion, each peptide $x$ is assigned a neutral monoisotopic mass $m(x)$, a theoretical fragment list $F(x)$, and a predicted spectrum $\phi(x)$. In the runs used for the manuscript, predicted spectra were obtained from Koina/Prosit with charge 2 and collision energy 27.

**Local candidate windows.** All comparisons are local in precursor-mass space. For a query peptide $q$, target and decoy competitors are restricted to

$$\mathcal{T}_q = \{t \in \mathcal{T} : |m(t) - m(q)| \leq \varepsilon_m\}, \tag{1}$$

$$\mathcal{D}_q = \{d \in \mathcal{D} : |m(d) - m(q)| \leq \varepsilon_m\}. \tag{2}$$

When $q$ itself is a target query, it is removed from $\mathcal{T}_q$ before computing the best false-target score. Likewise, when $q$ is a decoy query, it is removed from $\mathcal{D}_q$ before computing the best within-decoy competitor. In the default script configuration, $\varepsilon_m = 0.5$ Da.

**Similarity scores and nearest competitors.** The script computes predicted-spectrum cosine similarity,

$$s_{\cos}(q, x) = \cos\big(\phi(q), \phi(x)\big), \tag{3}$$

computed on sparse peak lists with the same fragment tolerance. In the default script configuration, $\varepsilon_f = 0.02$ Da.

For any chosen score $s$, the best false target and best decoy are

$$t^*(q) = \arg\max_{t \in \mathcal{T}_q} s(q, t), \tag{4}$$

$$d^*(q) = \arg\max_{d \in \mathcal{D}_q} s(q, d). \tag{5}$$

For target queries, $d^*(q)$ is the nearest effective decoy and therefore serves as the basic non-stealability diagnostic. The quantity plotted in the manuscript is typically the cosine distance to that nearest decoy,

$$\text{dist}_{\cos}(q, d^*(q)) = 1 - s_{\cos}(q, d^*(q)). \tag{6}$$

Small values indicate collision-prone target-decoy neighborhoods.

**Null competition and expected target-win rate.** For target and decoy queries alike, the script compares the best false-target score with the best decoy score and records the winner. Because the local candidate counts are not necessarily balanced, the expected target-win rate is not hard-coded to 0.5. Instead, for each query the script records

$$\mathbb{E}[\text{target win} \mid q] = \frac{|\mathcal{T}_q|}{|\mathcal{T}_q| + |\mathcal{D}_q|}, \tag{7}$$

with the query itself removed from its own class when appropriate. The observed winner proportions can then be compared against this locally expected rate rather than against an idealized global 50/50 split.

**Random null spectra.** To probe the null space more broadly, the script also simulates synthetic random spectra. The procedure is intentionally simple but keeps the precursor-mass and peak-count scales realistic:

1. Sample a precursor mass $m$ with replacement from the empirical masses of all target and decoy peptides in the effective library.
2. Sample a peak count $K$ with replacement from the empirical peak counts of the same library.
3. Draw $K$ fragment m/z values independently from a uniform distribution on $[100, \min(\max(300, m), 2200)]$ and sort them.
4. Draw $K$ intensities independently from an exponential distribution, then L2-normalize the resulting spectrum.
5. Search this synthetic spectrum against local target and decoy competitors in the same precursor window and record the best target and best decoy scores.

This null model does not aim to mimic real fragmentation chemistry. Its purpose is to create unstructured spectra with realistic precursor-mass and peak-count scales, providing a complementary stress test for local target/decoy balance.

**Reproducibility template.** A representative command used to generate the diagnostics is shown below.

```
PLOT_PDFS=1 python3 diagnostics.py \
  --fasta /path/to/library.fasta \
  --outdir /path/to/output \
  --workers 16 \
  --parallel-batch-size 512 \
  --koina-url https://koina.wilhelmlab.org \
  --koina-model Prosit_2020_intensity_HCD \
  --precursor-charge 2 \
  --collision-energy 27 \
  --koina-batch-size 1000 \
  --top-suspicious 100000 \
  --random-null-queries 1000000
```

The environment variable `PLOT_PDFS=1` switches the plot backend from PNG to PDF. The script also writes a `run_config.json` snapshot and cache tables that make it easy to reproduce the exact diagnostic run.

#### Note S4. Closest target–decoy pairs classification

To analyze the "target protection" property of different decoy generators, we identified the closest decoy for each target based on cosine distance and examined these pairs. Figure S4A shows the distribution of cosine similarity scores for the top 20,000 closest pairs derived from the yeast proteome. ESM2-based generators exhibit a more favorable score distribution, potentially reducing the number of spectral collisions between targets and decoys.

To further investigate the nature of high-similarity collisions, we classified the top 1,000 closest pairs for each generator (Figure S4B) into four categories:

- $I \leftrightarrow L$: the decoy differs from the target only by substitutions between isoleucine and leucine;
- $XY \leftrightarrow YX$: the decoy differs from the target by a single swap of adjacent amino acids;
- $I \leftrightarrow L, XY \leftrightarrow YX$: the decoy differs by both $I \leftrightarrow L$ substitutions and one adjacent amino acid swap;
- `rest`: all other cases.

For the `Reverse` and `Shuffle` generators, the majority of the top 1,000 closest pairs fall into the $I \leftrightarrow L$ and $XY \leftrightarrow YX$ categories, particularly for short peptides (Figure S4C–D). This highlights an increased likelihood of rare but challenging collisions within the limited sequence space of short peptides.

A substantial number of close target–decoy pairs falling into the `rest` category produced by `DIA-NN` generator reveals its inherent structure: multiple non-isobaric point mutations typically change the mass of the peptide, so in close target–decoy pair decoy is typically produced by a peptide that is very different from the target.

To further probe the structure of close target–decoy pairs and investigate the observed asymmetry of the `Reverse` and `Shuffle` generators between target and decoy queries (Figure S2B–C), we examined how often the closest decoy to a target corresponds to a decoy generated specifically for that target (Figure S5). For `Reverse` and `Shuffle`, this occurs in approximately 5–10% of cases. A similar proportion is observed for `DIA-NN`, which does not exhibit such asymmetry, suggesting that this factor alone does not explain the observed behavior. In contrast, ESM2-based generators do not produce such structured target–decoy pairs across peptide lengths.

#### Note S5. End-to-end search pipeline

**Pipeline overview.** The diagnostics were run with the `ms_pipeline` workflow, which orchestrates database search, optional rescoring, and post hoc calibration analysis. The pipeline accepts one or more spectrum files, a target FASTA, and a decoy-aware search configuration. In the experiments described here, the primary workflow was Sage, with optional Oktoberfest rescoring.

The pipeline first constructs the searched FASTA. In `entrapment-mode=none`, the provided FASTA is searched as-is. In `entrapment-mode=foreign`, a foreign-proteome FASTA is appended after prefixing its accessions with `ENTRAP_`. The code also supports a paired shuffled-peptide entrapment mode for additional experiments. Every run writes a `run.json` manifest containing input paths, hashes, and search configuration snapshots.

**Search and optional rescoring.** For Sage runs, the pipeline writes a Sage configuration file, launches the search, and then parses the resulting PSM table. If `run-oktoberfest` is supplied, the same search results are passed to Oktoberfest for rescoring. The analysis stage then produces both

PSM-level and peptide-level summaries. Peptide-level tables are obtained by taking the best q-value per peptide and carrying along a representative protein annotation from the best-scoring row.

Both Sage and Oktoberfest were executed with default settings except disabled decoy generation in Sage since we provide decoys as a part of the library.

**Entrapment labeling and FDP bounds.** Entrapment validation follows the logic described in the manuscript. Accepted rows are partitioned into original-target and entrapment-derived identifications using protein accessions. With the default `unambiguous` strategy implemented in the pipeline, a row is labeled as entrapment only if all associated proteins belong to the entrapment space; rows shared between original and entrapment proteins are marked ambiguous and excluded from both counts in the bound calculations.

For a q-value threshold $q$, let $N_T(q)$ be the number of accepted original-target identifications and $N_E(q)$ the number of accepted entrapment identifications. The lower and upper FDP estimates reported by the pipeline are

$$\mathrm{FDP}_{\mathrm{lower}}(q) = \frac{N_E(q)}{N_T(q) + N_E(q)}, \tag{8}$$

$$\mathrm{FDP}_{\mathrm{upper}}(q) = \frac{N_E(q)\,(1 + 1/r)}{N_T(q) + N_E(q)}, \tag{9}$$

where $r$ is the effective foreign-to-original search-space ratio that is estimated by digesting the target and entrapment FASTAs under the same enzyme settings used in the search.

**Outputs used in the manuscript.** The main outputs are identification-count curves versus q-value threshold and score-distribution plots. For Sage, the pipeline writes files such as:

- `sage_psm_counts_vs_q.csv`,
- `sage_psm_entrapment_bounds_vs_q.csv`, and
- their peptide-level counterparts.

Additional score-distribution plots are generated for the main score, matched-peaks count, and longest ion-series features, both overall and stratified by peptide length. Equivalent output families are generated for Oktoberfest when rescoring is enabled.

**Representative commands.** Representative command lines are shown below.

```
python3 -m ms_pipeline.ms_pipeline \
  --spectra /path/to/spectra.mzML \
  --target-fasta /path/to/library.fasta \
  --entrapment-mode none \
  --experiment-name some_name \
  --output-dir /path/to/output/dir \
  --sage-bin /path/to/sage
```

```
python3 -m ms_pipeline.ms_pipeline \
  --spectra /path/to/spectra.mzML \
  --target-fasta /path/to/library.fasta \
  --entrapment-mode foreign \
  --entrapment-prefix ENTRAP_ \
```

```
  --entrapment-fasta /path/to/entrapment/library.fasta \
  --experiment-name some_name \
  --output-dir /path/to/output/dir \
  --sage-bin /path/to/sage
```

```
# Oktoberfest rescoring can be enabled by supplying:
--run-oktoberfest
```

#### Note S6. End-to-end search benchmarks

Evaluation across multiple species (Figures S6, S7) reveals distinct patterns in decoy hardness. As expected, different decoy generators exhibit varying levels of "hardness" depending on the experimental setup. For instance, the `Reverse` generator appears "harder" than `Shuffle` for yeast data (Figure S6A–B), but becomes comparatively "easier" for human data (Figure S7A–B). Notably, in both datasets, ESM2-based decoys consistently become "easier" as the masking percentage increases (Figures S6E–I, S7E–I).

Results from Oktoberfest rescoring and comparisons between PSM- and peptide-level analyses show consistent trends (Figure S8). The number of identifications at both the PSM and peptide levels behaves similarly for a given decoy generator when applied to the same dataset. While Oktoberfest rescoring reduces the differences between decoy generators, it does not eliminate them entirely.

#### Note S7. Limitations and next steps

The diagnostics introduced in this work are intentionally designed to be largely search-engine independent. This provides conceptual clarity but also introduces an important limitation: Level 1 and Level 2 analyses can only approximate the behavior of a complete peptide identification pipeline. In particular, the spectral-space geometry derived from Prosit predictions does not capture every signal used by modern search engines, such as retention time, ion mobility, isotope structure, or nonlinear feature interactions used by rescoring models. Similarly, sequence-only separability primarily reveals generator fingerprints rather than search-engine calibration.

These limitations, however, highlight an important opportunity. The target–decoy framework naturally produces rich statistical signals that can be exploited not only for FDR estimation but also for diagnosing the behavior of search pipelines themselves. Deviations from the expected target–decoy balance—whether in nearest-neighbor structure, score distributions, or length-stratified collision patterns—can reveal situations in which search engines behave unexpectedly. In this sense, target–decoy statistics provide a practical lens for detecting potential failure modes, including shortcut learning, score miscalibration, or local target–decoy collisions that may compromise identification reliability.

An alternative direction for decoy generation is to consider adversarial learning frameworks, such as generative adversarial networks (GANs) [1], where a generator is trained to produce decoy peptides that are difficult to distinguish from targets under a discriminator model. Such approaches aim to better match the distribution of target sequences and may help mitigate construction-specific artifacts, thereby improving "null exchangeability" in a data-driven manner. At the same time, overly realistic decoys may increase the risk of violating "target protection", and therefore require careful regularization. Extensions based on sequential generative models, such as SeqGAN [2], are particularly relevant given the sequential structure of peptide data, and could provide a principled way to generate decoys that resemble the target. While we do not explore these approaches here, they represent a natural direction for future work.

Another limitation of this study is that we focus on DDA-style search pipelines, while data-independent acquisition (DIA) remains the dominant paradigm in many applications. From theoretical standpoint, the definition of "good" decoys is different for DDA and DIA since chimeric MS2 spectra in DIA are more tricky to handle from the theoretical standpoint. In practice, benchmarking alternative decoy generators in DIA is currently constrained by tooling. The most widely used engine, DIA-NN, does not expose an interface for supplying custom decoy libraries, and therefore effectively fixes the decoy generation strategy. While OpenSWATH in principle allows greater flexibility, integrating custom decoys requires substantial modification of the workflow and is not straightforward in routine use. As a result, we did not include DIA benchmarks here. Extending this analysis to DIA remains an important direction, as competitive structure and calibration may behave differently in library-based settings.

Future work could therefore expand the role of target–decoy analysis beyond generator benchmarking toward systematic search-engine diagnostics. For example, diagnostics could be integrated directly into benchmarking frameworks that compare search engines under controlled perturbations of the decoy database. Tunable generators, including PLM-based approaches, could be used to deliberately construct easy or adversarial decoy regimes in order to probe the robustness of scoring models. Another promising direction is the development of automated statistical tests that flag unusual target–decoy patterns during routine analyses, potentially serving as early-warning signals of calibration issues.

One more potential direction is to extend these diagnostics to more diverse search spaces. Although our study focuses primarily on conventional tryptic shotgun proteomics and HLA-I immunopeptidomics, other workflows including the use of alternative proteases, searches with variable post-translational modifications (PTMs), and open-search pipelines are becoming increasingly common. These settings induce different search-space structures, particularly in the case of open search. Nevertheless, we hypothesize that the diagnostics developed here should generalize to these regimes. For closed-search workflows involving alternative enzymes or variable modifications, this generalization is relatively straightforward. Open search, however, would require a more sophisticated Level 2 metric: rather than relying on a simple cosine distance between theoretical spectra, the distance measure should account for possible mass shifts at arbitrary peptide positions, maximizing spectral similarity under the mass-shift constraints imposed by the search engine.

Finally, the geometric analyses presented here could be extended to richer peptide representations that incorporate additional experimental dimensions such as retention time or ion mobility. Such extensions would bring the diagnostic space closer to the true scoring landscape used by modern identification pipelines.

### Supplementary References

[1] Ian J Goodfellow, Jean Pouget-Abadie, Mehdi Mirza, Bing Xu, David Warde-Farley, Sherjil Ozair, Aaron Courville, and Yoshua Bengio. Generative adversarial nets. *Advances in neural information processing systems*, 27, 2014.

[2] Lantao Yu, Weinan Zhang, Jun Wang, and Yong Yu. Seqgan: Sequence generative adversarial nets with policy gradient. In *Proceedings of the AAAI conference on artificial intelligence*, volume 31, 2017.
